# Monomodular *Pseudomonas aeruginosa* phage JG004 lysozyme (Pae87) contains a bacterial surface-active antimicrobial peptide-like region and a possible substrate-binding subdomain

**DOI:** 10.1101/2022.01.10.475687

**Authors:** Roberto Vázquez, Mateo Seoane-Blanco, Virginia Rivero-Buceta, Susana Ruiz, Mark J. van Raaij, Pedro García

## Abstract

Phage lysins are a source of novel antimicrobials to tackle the bacterial antibiotic resistance crisis. The engineering of phage lysins is being explored as a game-changing technological strategy for introducing a more precise approach in the way we apply antimicrobial therapy. Such engineering efforts will benefit from a better understanding of lysin structure and function. In this work, the antimicrobial activity of the endolysin from *Pseudomonas aeruginosa* phage JG004, termed Pae87, has been characterized. This lysin had been previously identified as an antimicrobial agent candidate, able to interact with the Gram-negative surface and disrupt it. Further evidence is hereby provided on this matter, based on a structural and biochemical study. A high-resolution crystal structure of Pae87 complexed with a peptidoglycan fragment showed a separate substrate-binding region within the catalytic domain, 18 Å away from the catalytic site and located at the opposite side of the lysin molecule. This substrate binding region was conserved among phylogenetically related lysins lacking an additional cell wall binding domain, but not among those containing such a module. Two glutamic acids were identified as relevant for the peptidoglycan degradation activity, although Pae87 antimicrobial activity was seemingly unrelated to it. In contrast, an antimicrobial peptide-like region within Pae87 C-terminus, named P87, was found to be able to actively disturb the outer membrane and have antibacterial activity by itself. Therefore, we propose an antimicrobial mechanism for Pae87 in which the P87 peptide plays the role of binding to the outer membrane and disrupting the cell wall function, either with or without the participation of Pae87 catalytic activity.

**Synopsis:** The structure of the monomodular *Pseudomonas aeruginosa* bacteriophage JG004 lysin Pae87 is presented and investigated in relationship with its function repurposed as an antimicrobial agent. The structure with its peptidoglycan ligand revealed a possible cell wall binding region. A C-terminal antimicrobial peptide-like region is shown to be important for disrupting the bacterial cell wall.

## 1. Introduction

Antibiotic resistance is becoming one of the most serious threats to public health worldwide since the number of multi-resistant bacterial strains is growing progressively. Therefore, the search for alternative treatments to standard antibiotics to fight these pathogenic ‘superbugs’ is becoming more urgent (O’Neill, 2016). Bacteriophage (phage) lysins are a promising strategy currently viewed as a feasible approach towards the development of novel, potentially marketable antibacterial agents (Abdelkader *et al*., 2019). Lysins are highly evolved enzymes produced by phages at the end of the lytic infection cycle to degrade the bacterial peptidoglycan, leading to cell lysis and phage progeny release. For about twenty years, phage lysins have been widely investigated as novel antimicrobial agents to treat bacterial infections caused, mainly, by Gram-positive pathogens. This mechanism relies on adding the purified enzyme exogenously (“lysis from without”) and provokes the rapid degradation of the substrate (the peptidoglycan) and, thus, the lysis and death of the susceptible bacteria, including multi-resistant strains. Lysins have demonstrated several advantages over standard antibiotics including: 1) rapid killing activity against both stationary-and exponential-phase bacteria, practically within a few minutes of contact with the peptidoglycan substrate; 2) effective against multi-drug-resistant bacteria; 3) specificity to the target pathogen, especially against Gram-positive bacteria, which allows the preservation of the normal microbiota; 4) resistance appearance seems very unlikely, probably due to the conservation of its substrate, the peptidoglycan; 5) synergistic effect with other lysins or antibiotics; 6) efficient lethal activity against colonizing pathogens growing on mucosal surfaces and/or in biofilms (Pastagia *et al*., 2013).

Lysin architecture varies depending on their origin, but those from phages infecting Gram-positive bacteria usually have a modular structure consisting of one or two catalytic domains, which harbor lytic activity against the host species, and a C-terminal cell wall binding domain (CWBD), which recognizes a cell wall trait specific to the bacteria that it targets. On the contrary, the great majority of lysins from phages infecting Gram-negative bacteria display a globular organization containing only one catalytic domain (Vázquez, García, *et al*., 2021). The catalytic domains are responsible for the cleavage of a specific bond within the peptidoglycan and based on the bond they break, lysins can be classified as glycosidases, N-acetyl-glucosamine amidases (NAM-amidases), or endopeptidases (Dams & Briers, 2019). Among glycosidase lysins, which are those that act within the glycan strand of N-acetyl-glucosamine (NAG) and N-acetylmuramic acid (MurNAc), two subclasses can be recognized. N-acetylmuramidases, just muramidases or lysozymes, hydrolyze the bond between MurNAc and NAG, at the reducing side of the former, while N-acetylglucosaminidases break the bond between NAG and MurNAc, at the reducing side of NAG. The CWBDs that many lysins bear are responsible for the specific recognition of the insoluble substrate and the high-affinity binding of these enzymes to the susceptible bacteria (Guillen *et al*., 2010; Low *et al*., 2011).

In a previous study (Vázquez, Blanco-Gañán, *et al*., 2021), the lysin from *Pseudomonas aeruginosa* phage JG004, termed Pae87, was mined from a dataset of phage lysin sequences (Vázquez, García, *et al*., 2021; Vázquez *et al*., 2020) on the basis of containing a putative C-terminal positively charged, antimicrobial peptide (AMP)-like region. The presence of such a region was probed under the hypothesis that it would enable Pae87 to disrupt the Gram-negative outer membrane (OM), often cited as a barrier to lysin activity from without (Briers & Lavigne, 2015). Pae87 was demonstrated to have an antimicrobial effect against *P. aeruginosa* and some other Gram-negative pathogens (Vázquez, Blanco-Gañán, *et al*., 2021). Nowadays, insightful knowledge of the structure of lysins, as well as implications for their function (either as exogenous antimicrobials or as lysis effectors for the virion particles release from the host cell) is a must to engage in protein-based antimicrobial engineering. An example is the combinatorial engineering of lysin modules or the derivation of AMPs from lysins (Duyvejonck *et al*., 2021; Thandar *et al*., 2016). Therefore, in this work, we provide a fine characterization of the structural elements that contribute to the observed activities of Pae87. We did so by (i) obtaining the crystal structure of the protein; (ii) examining putative catalytic residues by point mutation; and (iii) testing the antimicrobial and membrane permeabilizing activity of Pae87 and the AMP derived from its C-terminal region, named P87.

## 2. Materials and methods

### 2.1. Bacterial strains, media, and growth conditions

All Gram-negative bacteria used in this work (*P. aeruginosa, Escherichia coli, Acinetobacter baumannii, Acinetobacter pitti, Klebsiella pneumoniae*) were grown in Lysogeny Broth (LB, NZYTech) at 37°C with aeration (200 rpm shaking), except for *Moraxella catarrhalis*. This bacterial species, plus some Gram-positive ones (*Staphylococcus aureus, Streptococcus pyogenes*, and *Streptococcus* Milleri group strain) were cultured in Brain Heart Infusion broth (BHI, Condalab) at 37°C. *M. catarrhalis* and *S. aureus* were shaken at 200 rpm when grown in liquid culture. *Streptococcus pneumoniae* was grown in C medium adjusted at pH 8.0 (Lacks & Hotchkiss, 1960) supplemented with 0.08% yeast extract (C+Y) at 37°C without shaking. For solid cultures, all Gram-positives and *M. catarrhalis* were grown in blood agar plates, while Gram-negatives were grown in LB agar. Details on the bacterial strains used in this work can be found in (Vázquez, Blanco-Gañán, *et al*., 2021).

### 2.2. Plasmids, oligonucleotides, and overlap extension mutagenesis

Plasmids and oligonucleotides used throughout this work are included in Table S1. As explained in (Vázquez, Blanco-Gañán, *et al*., 2021), a synthetic gene encoding Pae87 (*pae87*) was obtained from GenScript, and cloned into a pET28a(+) vector in frame with an N-terminal 6×His tag. This expression plasmid, named pET-PA87, was heat shock-transformed into *E. coli* BL21(DE3). For the construction of *pae87* mutants (E29A, E46A, and E29A/E46A), an overlap extension PCR protocol was performed. Briefly, plasmid pET-PA87 was used as a template for two separate PCR reactions per mutation. Each of those reactions amplified a fragment of *pae87* gene in such a way that both fragments shared an overlapping section (≈ 20 nt) in which the mutated bases were located. A third PCR reaction was then performed using the *pae87* flanking primers pae87_f and pae87_3’ and a mixture of the resulting amplification products of the previous step as a template. Final PCR products were purified, digested with NdeI and HindIII, cloned into a pre-digested pET-28a(+) vector, and then transformed into *E. coli* DH10B. Colonies were screened by PCR and vectors putatively bearing the mutated gene were sequenced (Secugen, Centro de Investigaciones Biológicas Margarita Salas, Madrid, Spain). Final vectors were transformed into *E. coli* BL21(DE3) for expression.

### 2.3. Protein expression and purification

For the production of recombinant Pae87-based proteins, the appropriate strains were cultured in 1 l LB in the presence of 50 μg ml^-1^ kanamycin up to OD_600_ ≈ 0.6-0.8. Then, expression was induced with 0.4 mM isopropyl-β-D-thiogalactopyranoside and incubation was resumed at 20°C for up to 48 h. After centrifugation (12,000 × *g*, 20 min, 4°C), the pelleted biomass was resuspended in ≈ 30 ml of 20 mM sodium phosphate buffer (NaPiB), pH 7.4, containing 0.3 M NaCl and 40 mM imidazole. These cell suspensions were disrupted by sonication and cell debris were separated again by centrifugation (18,000 × *g*, 20 min, 4°C). The supernatants containing the protein extracts were then applied onto a HisTrap FF 5 ml column (GE Healthcare) loaded with nickel ions using an ÄKTA Start liquid chromatography machine (GE Healthcare). After a thorough washing step with the resuspension buffer, 20 mM NaPiB, pH 7.4, containing 0.3 M NaCl and 0.5 M imidazole was used to elute the purified protein. The buffer of the purified fractions was exchanged to 20 mM NaPiB, pH 7.4, containing 150 mM NaCl prior to assaying the proteins using HiTrap Desalting 5 ml columns. The concentration of purified proteins was estimated using the predicted molar extinction coefficients (Table S2) with A_280_ measurements. Protein samples were maintained at 4°C for up to a month without apparent signs of precipitation or loss of activity.

### 2.4. Synthesis and quantification of peptide P87

Peptide P87 (LNTFVRFIKINPAIHKALKSKNWAEFAKR) was synthesized and provided by GenScript as a freeze-dried powder. It was dissolved in distilled water and concentration was estimated by measuring A_280_, with a molar extinction coefficient of 5,500 M^-1^ cm^-1^, as predicted by ProtParam (Wilkins *et al*., 1999). Diluted peptide aliquots were kept at −20°C.

### 2.5. P. aeruginosa PAO1 peptidoglycan purification and muralytic activity assay

*P. aeruginosa* PAO1 peptidoglycan purification and dye-release muralytic activity assay were conducted essentially as explained in (Vázquez, Blanco-Gañán, *et al*., 2021). Briefly, *P. aeruginosa* PAO1 cells were cultured in 1 l LB broth until the OD_600_ reached 0.8-1.0. Then, the culture was centrifuged (4000 × *g*, 15 min, 4°C) and resuspended in 20 ml PBS. 80 ml of 5% SDS were added, boiling for 30 min with vigorous shaking. After overnight incubation at room temperature, the suspension was ultracentrifuged (100,000 × *g*, for 60 min at 20°C), and the pellet was resuspended in distilled water and subjected to dialysis against water for 24 to 72 h to wash out as much SDS as possible. Then, the samples were ultracentrifuged in the same conditions as before and washed again as many times as necessary to remove all SDS (typically 1-3 times more, until a thick foam layer was no longer formed upon resuspension). Then, RNA, DNA, and proteins were eliminated by successive treatments of RNAse, DNAse, and trypsin as explained in (Vázquez, Blanco-Gañán, *et al*., 2021). Finally, the peptidoglycan sacculi were ultracentrifuged again, the supernatant was removed and the pellet was dried for 24-48 h at 37°C to determine the dry weight yield of the process (typically, 12 mg l^-1^ of initial culture).

Purified sacculi were dyed by resuspending them in a freshly prepared 0.02 M Remazol Brilliant Blue (RBB) solution in 0.2 M NaOH. An incubation of about 6 h was conducted at 37°C with shaking and then overnight at 4°C. After staining, several ultracentrifugation and washing steps with distilled water were conducted until supernatants were clear (usually, 3-4 washing steps). The resuspension water volume of the final pellets was adjusted to an A_595_ ≈ 1.5. For the dye release assay, 100 μl of the RBB-stained sacculi were centrifuged (12,000 × *g*, 20 min, 20°C) and the supernatant was discarded. Then, the pelleted sacculi were resuspended in 100 μl of a solution of NaPiB, pH 6.0, containing 150 mM NaCl and the desired concentration of enzyme or just buffer for the control. The samples were incubated for 10 min at 37°C and reactions were stopped by incubating further 5 min at 95°C. Samples were then centrifuged (12,000 × *g*, 20 min, 20°C) and the A_595_ of supernatants was determined using a VersaMax multi-well plate spectrophotometer (Molecular Devices).

### 2.6. Antimicrobial activity assays

Antimicrobial activity assays were performed by incubating a resting bacterial cell suspension in NaPiB, pH 6.0, with 150 mM NaCl at 37°C together with the corresponding antibacterial protein or peptide (Vázquez, Blanco-Gañán, *et al*., 2021). Bacteria at the mid-to-late exponential phase, as evaluated by turbidimetry, were harvested by centrifugation (3000 × *g*, 10 min, 4°C). The pelleted cells were resuspended in half the volume of buffer and plated onto a 96-well plate (100 μl per well). 100 μl of the same buffer containing the desired concentration of the compound to be tested were then added and the plate was incubated at 37°C for 2 h. Several measurements were performed on the treated resting cells, namely: OD_600_ monitoring, viable cell counts by plating 10-fold serial dilutions at the end of the experiment, and observation under a fluorescence microscope (Leica DM4000B with an HC PL APO 100×/1.40 oil objective and L5 [bandpass 480/40] and N2.1 [515/60] filters) stained with the BacLight LIVE/DEAD bacterial viability kit L-13152 (Invitrogen-Molecular Probes, containing SYTO9 and propidium iodide).

### 2.7. Analysis of degradation products

To analyze the degradation products that resulted from Pae87 activity, a similar protocol to that described in (Alvarez *et al*., 2016) was followed. 100 μl of the *P. aeruginosa* PAO1 purified peptidoglycan were centrifuged (12,000 × *g*, 20 min, room temperature) and resuspended again in the same volume of a suitable reaction buffer (20 mM NaPiB, pH 6.5, 100 mM NaCl, for Pae87 or 50 mM NaPiB, pH 4.9, for the positive control cellosyl). Then, 10 μg of the corresponding enzyme (or an equivalent volume of water for the negative control) were added and samples were incubated overnight at 37°C. Reactions were stopped by incubating at 98°C for 5 min. The tubes were then centrifuged (12,000 × *g*, 20 min, room temperature) and the supernatants were collected. 0.5 M borate buffer, pH 9.0 was added to adjust the pH of the samples to 8.5-9.0, and 10 μl of freshly prepared 2 M NaBH_4_ were added to reduce the sample at room temperature for 30 min. Next, pH was adjusted to 2.0-4.0 with 25% orthophosphoric acid. Then, the soluble muropeptides of each sample were separated by reverse-phase high-performance liquid chromatography (RP-HPLC) on a Kinetex C18 Column (1.7 μm, 100 Å, 150 × 2.1 mm, Phenomenex) coupled to an LXQ mass spectrometer (MS) equipped with a linear ion trap (Finnigan TM LXQTM, Thermo Scientific). Ionization was achieved by electrospray. Muropeptides were eluted at a flow rate of 0.4 ml min^-1^ with the following elution gradient: t = 0 min, 95% A; t = 0.5 min, 93% A; t = 3 min, 82% A; t = 11 min, 50% A; t = 12 min, 50% A; t = 12.1 min, 95% A, t = 15 min, 95% A; being A: 0.1% formic acid in water and B: 0.1% formic acid in 40% acetonitrile. The MS was operated on a double play mode in which the instrument was set to acquire a full MS scan (150–2,000 m/z) and a collision-induced dissociation (CID) spectrum on selected ions from the full MS scan. Spectra were analyzed using Xcalibur software (Thermo Scientific).

### 2.8. Pae87 production and crystallization

*E. coli* BL21(DE3) transformed with pET-PA87 was grown at 37ºC as described in section 2.3. After reaching an OD_600_ of about 0.6, cultures were cooled and Pae87 expression was induced with 0.5 mM of isopropyl-β-D-thiogalactopyranoside at 20ºC overnight. The cells were centrifuged at 6,000 × *g* for 10 min at 4ºC and pellets were stored frozen. Later, pellets from 1 l of culture were resuspended with 10 ml of lysis buffer (20 mM Tris pH 7.5, 0.5 M NaCl, 20 mM imidazole and 5% glycerol). Then, cells were disrupted with a Digital Sonifier 250 (Branson) and centrifuged at 15,000 × *g* for 45 min at 4ºC. Supernatants from the centrifugation of the disrupted cells were incubated with 2 ml of nickel-nitrilotriacetic acid (Ni-NTA) agarose (Jena Bioscience) on ice for 30 min. The mixture was then poured into an Econo-Pac Chromatography Column (Bio-Rad) and eluted by gravity. After a two column-volume wash with lysis buffer, the protein was eluted by passing 2 ml of elution buffer (20 mM Tris pH 7.5, 0.5 M NaCl, 0.5 mM imidazole and 5% glycerol) six times. After analysis by denaturing gel electrophoresis, the protein-containing fractions were dialyzed in a cellulose membrane tube (14 kDa MWCO, Merck-Millipore), first, against 1 l of 20 mM Tris-HCl, pH 7.5 and 0.2 M NaCl for 3-4 h at 4ºC and then, against 1 l of 20 mM Tris-HCl, pH 7.5 overnight at 4ºC. Prior to ion exchange chromatography, the dialyzed sample was centrifuged at 15,000 × *g* for 10 min at 4ºC to remove aggregates. Chromatography was performed using an ÄKTApurifier 10 FPLC system (Cytiva) with a RESOURCE™ Q 6 ml column. Proteins were eluted with a gradient of 0 to 1 M NaCl buffer with 20 mM Tris-HCl, pH 7.5. Pae87 eluted at approximately 0.18 M NaCl. Fractions containing pure Pae87 were desalted and concentrated with Amicon Ultra centrifugal filters of 3 kDa or 10 kDa MWCO (Millipore). Concentrated samples were centrifuged at 15,000 × *g* for 10 min prior to crystallization trials; the pellet was discarded. Pae87 concentration was estimated by measuring A_280_, using the predicted molar extinction coefficient (Table S2).

Proteins were crystallized using the sitting-drop vapor-diffusion technique in MCR crystallization plates (SWISSCI). Reservoirs were filled with 50 μl of different crystallization solutions. Drops contained 1 μl of protein (in 20 mM Tris-HCl, pH 7.5) plus 0.5 μl of crystallization solution. The best apo-protein crystal was obtained at 11 mg ml^-1^ Pae87, when 20% (w/v) polyethyleneglycol 8,000 and 0.1 M CHES-NaOH, pH 9.5 was used as crystallization solution. For the ligand-protein complex crystallization assay, 3 mg of the dried *P. aeruginosa* PAO1 peptidoglycan was solubilized by adding 120 μl of the purified Pae7 protein at 11 mg ml^-1^ in 20 mM Tris-HCl, pH 7.5. The mixture was centrifuged at 15,000 × *g* for 10 min at 4ºC to remove aggregates. Then, the supernatant was used in crystallization trials. The best crystal appeared when 0.1 mM Tris-HCl, pH 8.0, 20% (w/v) polyethyleneglycol monomethyl ether 5,000, 8% (v/v) ethylene glycol was used as the crystallization solution.

### 2.9. Crystallographic data collection, structure determination and analysis

Crystals were diffracted at the XALOC beamline of the ALBA-CELLS synchrotron (Cerdanyola del Vallès, Spain) (Juanhuix *et al*., 2014), using a Dectris Pilatus 6M detector. Diffraction images were processed inside the CCP4 suite (Winn *et al*., 2011). XIA2/DIALS (Winter, 2010) was used to index and integrate the images, POINTLESS (Evans, 2006) for space group determination and AIMLESS (Evans & Murshudov, 2013) for scaling. The crystal structure of the apo form was determined by molecular replacement with MOLREP (Vagin & Teplyakov, 2010), using PDB entry 5NM7 as a search model (Maciejewska *et al*., 2017). The protein model was modified using the graphics program COOT (Emsley *et al*., 2010) and refined with REFMAC5 (Murshudov *et al*., 2011). The protein-ligand complex structure was similarly obtained by molecular replacement, using molecule A of the apo form model as a search model. The resolution limit for data to be included was determined by paired refinement with PDB_REDO (Joosten *et al*., 2014). Models were validated with MolProbity (http://molprobity.biochem.duke.edu/) (Williams *et al*., 2018). PyMOL (The PyMOL Molecular Graphics System, Version 1.8 Schrödinger, LLC) was used for analysis and visualization of the models. Structure comparison was performed in the DALI server (Holm, 2020) and oligomerization parameters (accessible and buried surfaces, estimated dissociation energies) were analyzed with QtPISA (Krissinel, 2015).

### 2.10. Bioinformatic analyses

The lysin sequence data set published in (Vázquez, García, *et al*., 2021) and available at (Vázquez *et al*., 2020) was used as a sample for different analyses. Multiple sequence alignments were performed using Clustal Omega as implemented at the EMBL-EBI (Madeira *et al*., 2019) and visualized using JalView (Waterhouse *et al*., 2009). HeliQuest (Gautier *et al*., 2008), and EMBOSS charge (Rice *et al*., 2000) were used to predict physicochemical properties of protein helices and local charge plots amino acid sequences. To specifically determine the sequence of the AMP-like fragment on the C-terminus of Pae87, a score variable was calculated consisting in the half-sum of the min-max standardized net charge (NC) and hydrophobic moment (HM) for each 11-amino acid fragment of Pae87 (**Equation 1**):

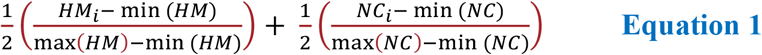

where HM_i_ and NC_i_ are specific values of HM or net charge, respectively, and HM and NC the respective sets of values for each variable. In this way, the score measured the magnitude of both parameters with equal weight and on a 0-1 scale.

Sequence Similarity Networks (SSNs) were generated for visually assessing the similarity clustering of sequence sets. For this purpose, the Enzyme Similarity Tool from the Enzyme Function Initiative server (EFI-EST) was employed (Zallot *et al*., 2019). Briefly, this tool performs a local alignment from which every possible pair of sequences receives a score similar to the *E*-value obtained from a typical BLAST analysis. A threshold score value was selected for the SSN so that below such threshold, sequence pairs were considered non-similar and, therefore, the pair would not be connected in the resulting representation. The score was selected so that sequence pairs whose similarity was below 30-40% were deemed non-similar. The SSN graphs were produced using Cytoscape 3 with identity %-based Prefuse Force Directed layout (Shannon *et al*., 2003).

### 2.11. Circular dichroism

Circular dichroism spectra were acquired at 4°C in 20 mM NaPiB, pH 6.5, 150 mM NaCl, using a Jasco J700 spectropolarimeter equipped with a temperature-controlled holder. Far UV spectra were recorded from 260 to 200 nm in a 1 mm path length quartz cuvette. Each spectrum was obtained by averaging 5 accumulations collected at a scan rate of 50 nm min^-1^ and 2 s of response time. Buffer spectra were subtracted from 0.1 mg ml^-1^ peptide spectra and molar ellipticity per residue was calculated. Far UV spectra collected with different concentrations of 2,2,2-trifluoroethanol (TFE) were used to predict the ability of peptides to form secondary structures in the presence of biological membranes.

### 2.12. Outer membrane permeabilization assay

The physiological effects of the interaction of proteins or peptides with the *P. aeruginosa* PAO1 OM were assayed by using N-phenyl-1-naphthylamine (NPN) as a fluorescent probe (Loh *et al*., 1984; Helander & Mattila-Sandholm, 2000). The assay was conducted with suspensions of PAO1 resting cells prepared from an actively growing culture at ≈ 10^8^ CFU/ml pelleted (3,000 × *g*, 10 min, 20°C) and resuspended in half the volume of 20 mM NaPiB, pH 6.0, 150 mM NaCl, 100 mM sorbitol. The sorbitol was added as an additional osmoprotectant to maintain cellular integrity. 100 μl of this suspension were added to each well of a FluoroNunc 96-well plate together with 50 μl of 40 μM NPN and 50 μl of the enzyme or peptide at the corresponding concentration, or just buffer for the negative control. Fluorescence was then immediately recorded in a Varioskan Flash microplate reader (Thermo Scientific) with an excitation wavelength of 350 nm and 420 nm for fluorescence detection. Non-dyed PAO1 cells treated with just buffer were also incorporated as background measurement, and EDTA was used as a positive control for OM permeabilization. Samples were also observed by fluorescence microscopy using an A filter cube (Leica), with a bandpass of 340-380.

### 2.13. Fluorochrome labelling of Pae87

An N-hydroxysuccinimidyl (NHS) ester of Alexa488 fluorochrome (NHS-Alexa488, ThermoFisher) was used to fluorescently label purified Pae87. A labelling reaction was set up by incubating the dye together with the protein at a 1:5 (protein:dye) ratio in 20 mM NaPiB, pH 7.4, 100 mM NaCl, for 1 h at room temperature. At this pH, N-terminal amino groups would be more reactive than lateral chains amino groups, and therefore the dye would preferentially be incorporated at N-terminal. The reaction was stopped by adding 10% volume of 1 M Tris-HCl, pH 7.4. Free dye was then separated using a HiTrap Desalting FF 5 ml column on an ÄKTA Start liquid chromatography system with 20 mM NaPiB, pH 7.4, 100 mM NaCl, as the mobile phase. The much smaller molecule of the dye was longer retained within the column, while the dyed protein eluted earlier. The degree of labelling of each elution sample was calculated by estimating the protein and dye concentrations measuring, respectively, A_280_ and A_495_ and applying the Lambert-Beer law with the respective molar extinction coefficients. A_280_ was corrected with a factor of 0.11 (corresponding to A_280_/A_495_ for the dye). Only samples with a labelling degree of ≈ 1 were used for experiments.

### 2.14. Statistical analyses

Statistical analyses were performed either in R or using GraphPad InStat. Unless otherwise stated, quantitative differences between experimental conditions were analyzed to determine whether such differences were statistically significant under the assumption of a signification level α = 0.05 by using ANOVA. The *post hoc* tests of Tukey or Dunnett were further used for multiple comparisons. Unless otherwise stated, all results presented are mean ± standard deviation of at least three independent replicates.

## 3. Results

### 3.1. General analysis of the *Muramidase* family

Pae87 contains a single predicted catalytic domain that occupies most of its full length (90.3% coverage) and belongs to the recently described *Muramidase* family (PF11860, Pfam E-value 1.1×10^−63^). The *Muramidase* family is thought to comprise N-acetylmuramidases (EC 3.2.1.17) that, as predicted by Pfam, usually appear either among bacteria (mainly Proteobacteria) or bacteriophages from the Caudovirales family (Fig. 1*a*). Lytic enzymes containing a *Muramidase catalytic domain* comprise some architectures that range from a single *Muramidase catalytic domain* to *Muramidase* domains accompanied by different known CWBDs either at the N-or C-terminus (Fig. 1*b*). The SSN constructed with the set of *Muramidase* family representatives (Fig. 1*c*) contained in InterPro displays some similarity-based clustered groupings, such as that of *Pseudomonas*-contained proteins, or some Alphaproteobacteria clusters, but, in general, it does not provide evidence of clear taxonomically relevant subfamilies. In a previously curated collection of phage lysin sequences named 𝕊^*LYS*^ (Vázquez *et al*., 2020), *Muramidase* domains accounted for 1.56% of the total predicted domains (Fig. 1*d*), mostly represented in phages that infect Gram-negative bacteria (Fig. 1*e*), among which the *Muramidase* family ranked as one of the most common catalytic domains. In the subset of 𝕊^*LYS*^ that comprises just those entries bearing at least a *Muramidase catalytic domain*, hereinafter termed 𝕊^*MUR*^, the length distribution contains two subpopulations (Fig. 1*f*) that can be related to the presence or absence of a CWBD, usually located at the N-terminus (Fig. 1*gh*). 54.3% of the lysins in 𝕊^*MUR*^ contained two predicted domains (*i*.*e*., bear both a *Muramidase catalytic domain* and a CWBD) while the rest are thought to be mono-modular, as Pae87 itself. The SSN of 𝕊^*LYS*^ did not display a relevant taxonomical clustering (Fig. 1*h*), although the sample size was on the low side to be able to draw generalized conclusions.

**Figure 1.**
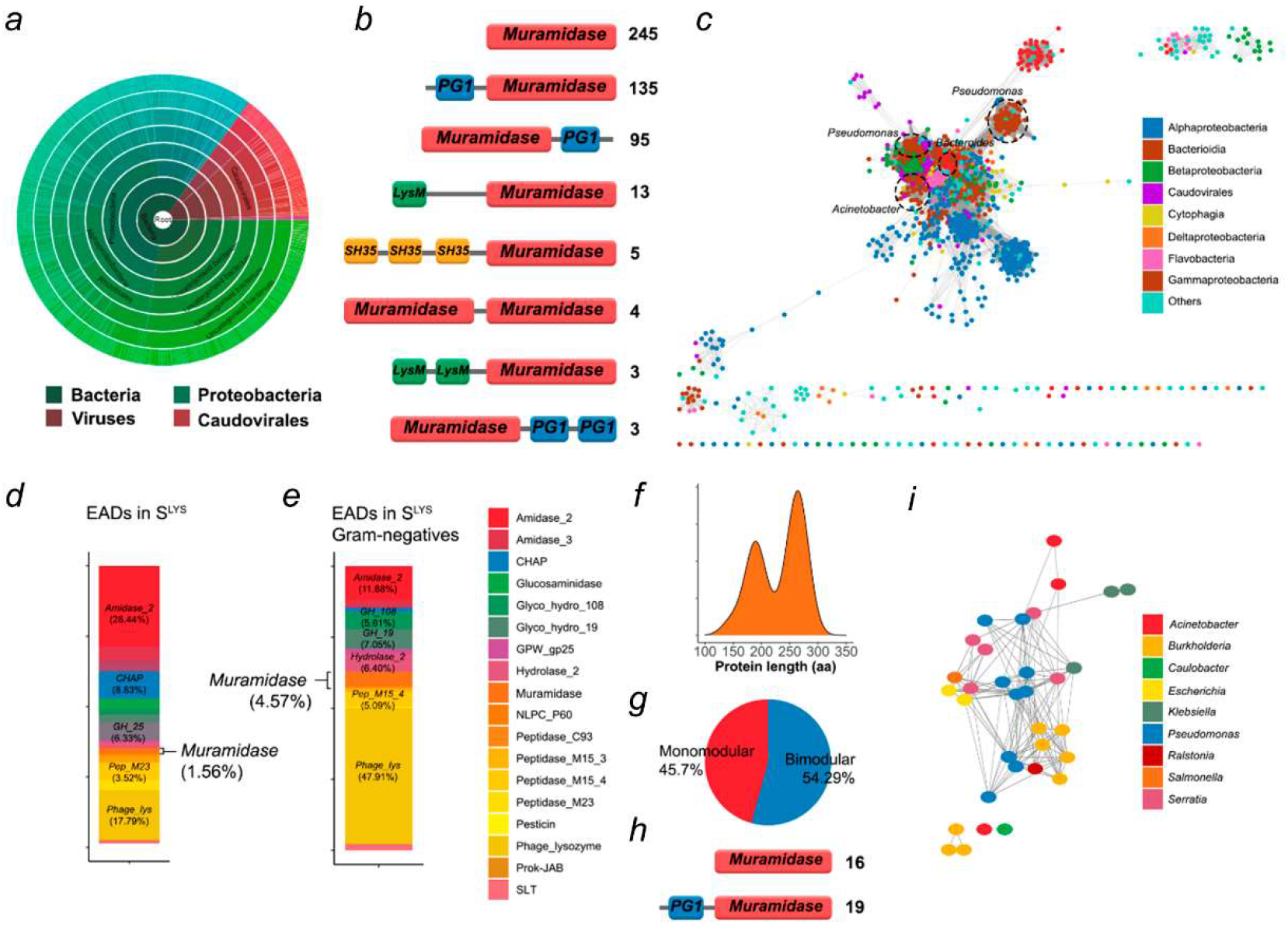
Overview of the *Muramidase* domain family. (*a*) Sunburst taxonomical representation of the organisms bearing predicted *Muramidase* (PF11860)-containing proteins as taken from Pfam. (*b*) Main architectures found among *Muramidase*-containing proteins in Pfam. The leftmost numbers are the number of representatives predicted to have such architecture (PG1 = *PG_binding_1* [PF01471], SH35 = *SH3_5* [PF08460], *LysM* = PF01476). (*c*) SSN comprising the predicted *Muramidase*-bearing proteins in InterPro IPR024408 entry, corresponding to *Muramidase* family. (*d, e*) Distribution of catalytic domains in phage lysin curated database 𝕊^*LYS*^ (*d*) or in the subset of 𝕊^*LYS*^ comprising only those lysins from phages that infect Gram-negative bacteria (*e*). (*f*) Length distribution of the *Muramidase*-containing lysins in 𝕊^*LYS*^ (𝕊^*MUR*^). (*g*) Distribution of the number of predicted domains in 𝕊^*MUR*^. (*h*) Architectures found in 𝕊^*MUR*^. (*i*) SSN of 𝕊^*MUR*^.

### 3.2. Three-dimensional structure of Pae87 and substrate binding region

While evidence has been previously provided for the antimicrobial activity of Pae87 (Vázquez, Blanco-Gañán, *et al*., 2021), to further elucidate the mechanism by which Pae87 interacts with susceptible bacterial cells, we determined crystal structures of the protein with and without a bound peptidoglycan fragment.

The protein without and with PAO1 peptidoglycan was crystallized as described in the methods section. The best apo-protein crystal diffracted only to limited resolution, but the best peptidoglycan-bound protein crystal diffracted X-rays to high resolution (Table 1). In the crystal packing, the peptidoglycan fragment contacts a neighboring protein molecule; this may be the reason why the ligand-bound form crystallized in different conditions and in a different crystal form that diffracted X-rays to higher resolution. A structure prediction search in HHpred (Zimmermann *et al*., 2018) showed that AP3gp15, a *Burkholderia* AP3 phage endolysin that belongs to the *Muramidase* PF family, had the most similar sequence to Pae87 (50% identity with 183 amino acids in the alignment). Therefore, the apo-protein was solved by molecular replacement, using the protein AP3gp15 [(Maciejewska *et al*., 2017) PDB entry 5NM7] as a search model.

**Table 1.** Crystallographic statistics of the Pae87 datasets.

|  | Pae87 | Pae87-peptidoglycan |
| --- | --- | --- |
| **Data collection** |  |  |
| Wavelength (Å) | 0.97926 | 0.97933 |
| Crystal-to-detector distance (mm) | 488.7 | 212.5 |
| Space group | *P*2<sub>1</sub>2<sub>1</sub>2<sub>1</sub> | *P*2<sub>1</sub>2<sub>1</sub>2<sub>1</sub> |
| Cell edges (a, b, c; Å) | 59.25, 68.09, 93.10 | 43.58, 61.17, 69.21 |
| Resolution range (Å) | 59.25-2.50 (2.60-2.50) | 69.21-1.27 (1.29-1.27) |
| Total number of reflections | 78046 (8657) | 314568 (14575) |
| Number of unique reflections | 13640 (1509) | 49510 (2412) |
| Completeness (%) | 99.7 (99.1) | 99.9 (100.0) |
| Multiplicity | 5.7 (5.7) | 6.4 (6.0) |
| CC 1/2<sup>a</sup> | 0.991 (0.860) | 1.000 (0.903) |
| R<sub>meas</sub> (all I+ & I-)<sup>b</sup> | 0.202 (1.087) | 0.046 (0.828) |
| <I/σ(I)> | 6.0 (2.1) | 17.6 (2.0) |
| Wilson B | 31.8 | 13.9 |
| **Refinement** |  |  |
| Resolution range (Å) | 55.02-2.50 | 45.85-1.27 |
| Reflections used | 12941 | 46988 |
| Reflections used for R-free | 658 | 2454 |
| R-factor/R-free<sup>c</sup> | 0.222/0.278 | 0.130/0.161 |
| **Model statistics** |  |  |
| Atoms (protein / peptidoglycan / CHES / PEG / ethylene glycol / water) | 2964 / 0 / 26 / 13 / 0 / 99 | 1506 / 50 / 0 / 14 / 52 / 121 |
| Average temperature factor (Å<sup>2</sup>, protein / peptidoglycan / CHES / PEG / ethylene glycol / water) | 60.3 / - / 89.0 / 73.4 / - / 41.4 | 20.7 / 20.9 / - / 42.2 / 44.4 / 35.1 |
| Ramachandran<sup>d</sup> (% favored / allowed) | 96.5 / 99.7 | 97.8/100.0 |
| RMSD (bonds, Å / angles, °)<sup>e</sup> | 0.0016 / 1.1 | 0.0073 / 1.4 |
| PDB code | 7Q4S | 7Q4T |

\*The values in parentheses correspond to the high-resolution shell
<sup>a</sup> CC1/2 correlation coefficient between intensity estimates from half data sets (Karplus & Diederichs, 2015).
<sup>b</sup> $R_{meas} = (\sum_{hkl} [n/n-1])^{1/2} \sum_i |I_{i,hkl}| \sum_i I_i(hkl)$
<sup>c</sup> $R = (\sum_{hkl} |F_{obs}(hkl) - F_{calc}(hkl)|) / \sum_{hkl} |F_{obs}(hkl)|$
<sup>d</sup> Calculated by MolProbity (Williams *et al.*, 2018)
<sup>e</sup> Calculated by REFMAC5 (Evans & Murshudov, 2013)

The crystallographic models of Pae87 comprise all 186 residues of the protein, plus up to two residues from the N-terminal purification tag. The apo-Pae87 and Pae87-peptidoglycan have two and one molecules in the asymmetric unit, respectively. Molecule B of the apo structure is less well-ordered than molecule A as evidenced by the average temperature factors (68.2 vs. 52.4 Å^2^, respectively). The root mean square deviations (RMSD) between molecules A and B of the apo-Pae87 structure and between molecules A or B of the apo structure and molecule A of the Pae87-peptidoglycan structure are 0.5 Å when Cα atoms are superposed. Since the structures are almost identical, we mainly describe the Pae87-peptidoglycan complex structure and only add a few details specific to the apo-Pae87 structure.

The crystallographic structure of Pae87 is almost identical to the C-terminal catalytic domain of the endolysin AP3gp15 (RMSD value of 1.1 Å on 183 superposed Cα atoms). Pae87 lacks the N-terminal CWBD present in AP3gp15 (Fig. 2*a*). The Pae87 structure shows the typical lysozyme-like α/β fold subdivided into two subdomains (Wohlkonig *et al*., 2010). The α-lobe is composed of α-helices 1 and 2 (Glu5-Glu29) and α6-12 (Gly109-Lys186). In the α-lobe subdomain, α2 is surrounded by the other α-helices. The other subdomain is traditionally known as the β-lobe because it contained a β-sheet in the first known endolysin structures (as it does in Pae87). In Pae87, the β-lobe is formed by β-strand 1 (Ile43-Glu46), α-helices 3, 4 and 5 (Arg47-Ser99), and β-strands 2 and 3 (Ala 100-Met108). The three β-strands form a small antiparallel β-sheet (Fig. 2*b*).

**Figure 2.**
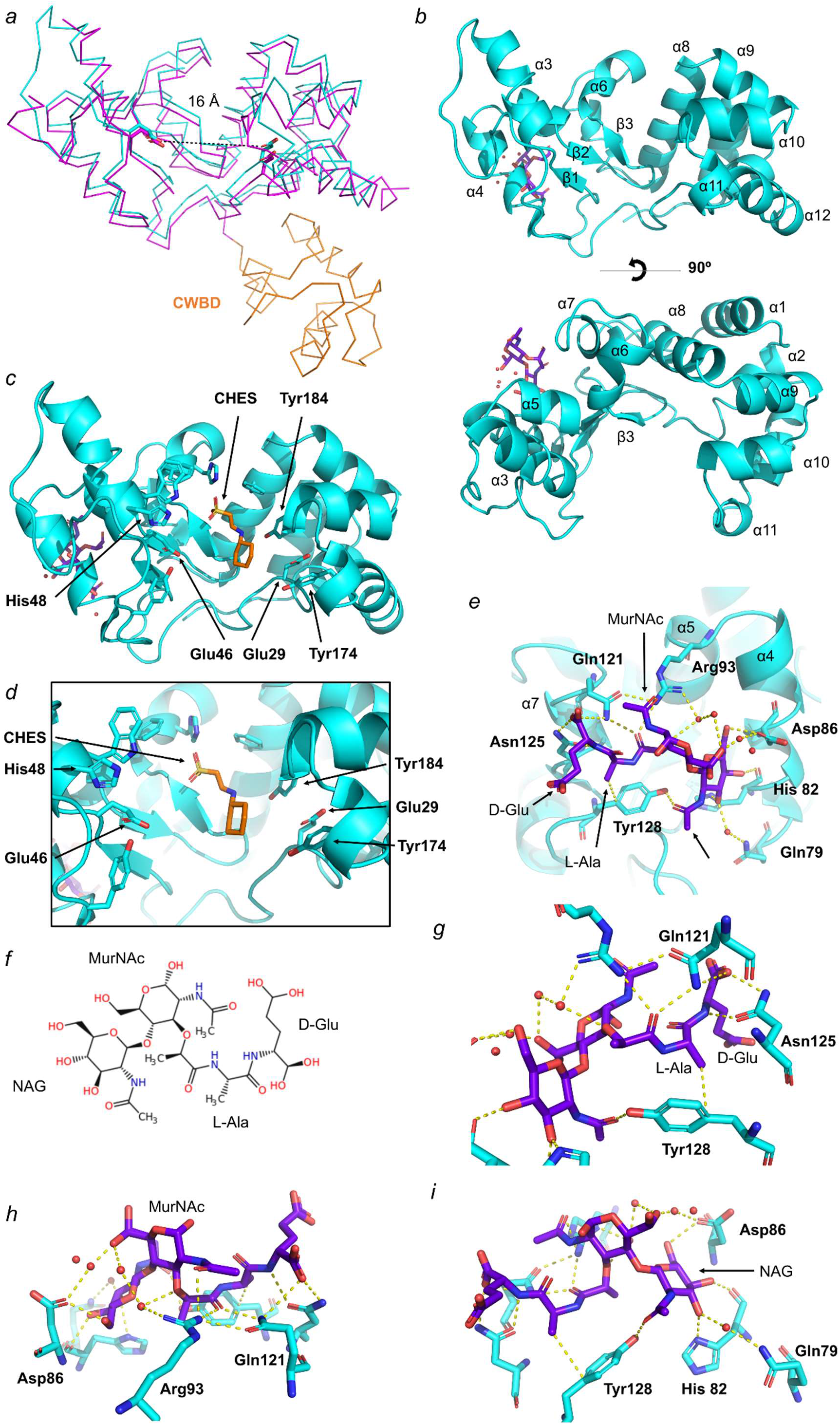
Pae87 structure. (*a*) Superimposition of the catalytic domains of Pae87 (cyan) on AP3gp15 (magenta), with the AP3gp15 CWBD shown in orange. Glu29 and Glu66 from Pae87 and Glu101 and Glu118 from AP3g015 are shown in stick representation. (*b*) Side view (above) and upper view (below) of the Pae87 protein model in ribbon representation (cyan) in front of the peptidoglycan fragment colored violet in stick representation. (*c*) View of a CHES molecule present in the apo-Pae87 model (orange, stick representation) and modelled in the putative catalytic site of the Pae87-peptiddoglycan structure. It is flanked by the putative catalytic amino acids (Glu29 and Glu46) and several aromatic residues in stick representation. (*d*) Close-up view of the active site depicted as in (*c*). (*e*) View of the peptidoglycan fragment depicted as in (*b*) with the binding amino acids (cyan, stick representation), the water molecules (red spheres) taking part in the hydrogen bond network (yellow dashed lines). (*f*) Schematic representation in Fischer projection of the peptidoglycan fragment bound to Pae87. (*g-i*) Close-up views of the peptidoglycan fragment components (D-Gly and L-Ala, MurNAc and NAG, respectively) depicted as in (*f*).

When Pae87 was crystallized together with the hydrolyzed peptidoglycan sacculi, we expected to see a fragment of it bound to the catalytic cleft. Surprisingly, the peptidoglycan fragment (comprised of linked NAG, MurNAc, L-Alanine and D-Glutamic acid units) was bound to α-helices 4, 5 and 7, on the opposite side to the putative catalytic pocket (Fig. 2*d*). This fragment was bound mainly by a hydrogen bond network formed by Gln79, His 82 and Asp86 (α4); Arg93 (α5); Gln121, Asn125 and Tyr128 (α7) (Fig. 2*g-i*). Tyr128 probably establishes a CH-π interaction with the L-Ala methyl group. The most important binding residues would be Arg93, Gln121, Asn125 and Tyr128, because they each have more than one bond with the ligand.

A multiple sequence alignment (MSA) analysis of the amino acids composing the peptidoglycan binding site using 𝕊^*MUR*^ (but removing two low-coverage examples, YP_009639957.1 and YP_337984) revealed that the residues relevant for the peptidoglycan binding were much more conserved in those lysins bearing a single *Muramidase* catalytic domain than in those which had an additional CWBD (Fig. 3*a*). Specifically, in the case of Tyr128 and Arg93, together with Gln121 and Asn125, both the amino acid frequency and the relative BLOSUM62 score (which is a similarity metric) were close to the maximum value, 1.0, for the non-CWBD bearing entries (Fig. 3*bc*). For the bimodular lysins of the *Muramidase* family, the scores were below 0.4 and 0.2 for Tyr128 and Arg93, respectively. The SSN in Fig. 3*d* shows that no sequence identity bias was introduced by the classification since the average identity percentage both within each subgroup and in the whole collection of sequences was around 46-52% and there was no apparent clustering.

**Figure 3.**
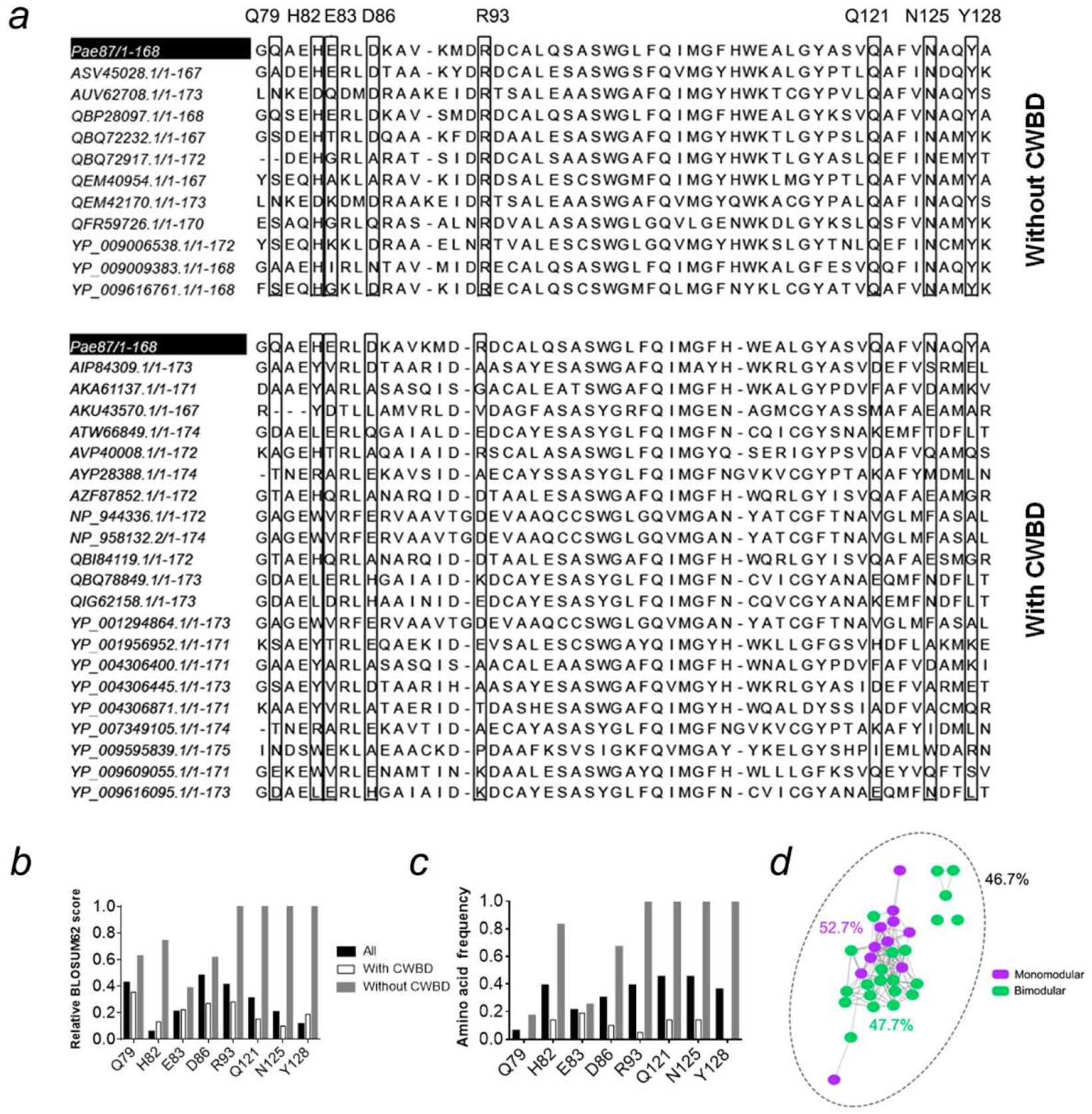
MSAs of the peptidoglycan binding site of lysins in 𝕊^*MUR*^. (*a*) MSA of the peptidoglycan binding site of *Muramidase* family lysins grouped by the presence or absence of an N-terminal CWBD. Residue coordinates indicated at the top are those of Pae87. (*b, c*) Residue conservation metrics for the residues putatively involved in contacts with peptidoglycan at the peptidoglycan binding site according to the Pae87 3D model (*b*: relative BLOSUM62 score; *c*: relative frequency of the Pae87 residue across the MSAs in each respective position). (*d*) SSN of the 33 sequences of the MSAs distinguished by the presence or absence of a CWBD. Percentages are average identities for each group. The entries AKU43570.1 and YP_009595839.1 were reclassified as non-CWBD bearing lysins since they contained an unidentified N-terminal end that could probably be a still undescribed CWBD.

### 3.3. Analysis of Pae87 catalytic center

A single glutamic acid has been pointed out to be involved in catalysis for AP3gp15 (Maciejewska *et al*., 2017). A corresponding Glu residue is also conserved in the Pae87 structure (Glu29, Fig. 4) and all of the 32 *Muramidase* sequence examples in the curated 𝕊^*MUR*^. Glu29 is located within a deep cleft between the subdomains of Pae87 structure (Fig. 2*b*), and thus this is considered to be the catalytic site. This glutamic acid residue is conserved in many endolysins, such as AP3gp15 (PDB entry 5NM7) (Maciejewska *et al*., 2017), Hen Egg-White Lysozyme (HEWL, PDB entry 4HPI) (Ogata *et al*., 2013) and the peptidoglycan hydrolase Auto (PDB 3FI7) (Bublitz *et al*., 2009). Most lysozymes have a catalytic site formed by two residues: a general acid and a general base – normally acidic amino acids (Glu or Asp). The C-terminal region of the central α-helix (α2) of the lysozyme-like α/β fold typically contains a conserved catalytic glutamate, but the general base is not well conserved and does not exist in some lysozymes. In HEWL, it is located 5 Å away in the other side of the cleft; and in the peptidoglycan hydrolase Auto, the general base is 14 Å away, in the antiparallel β-sheet (Bublitz *et al*., 2009).

**Figure 4.**
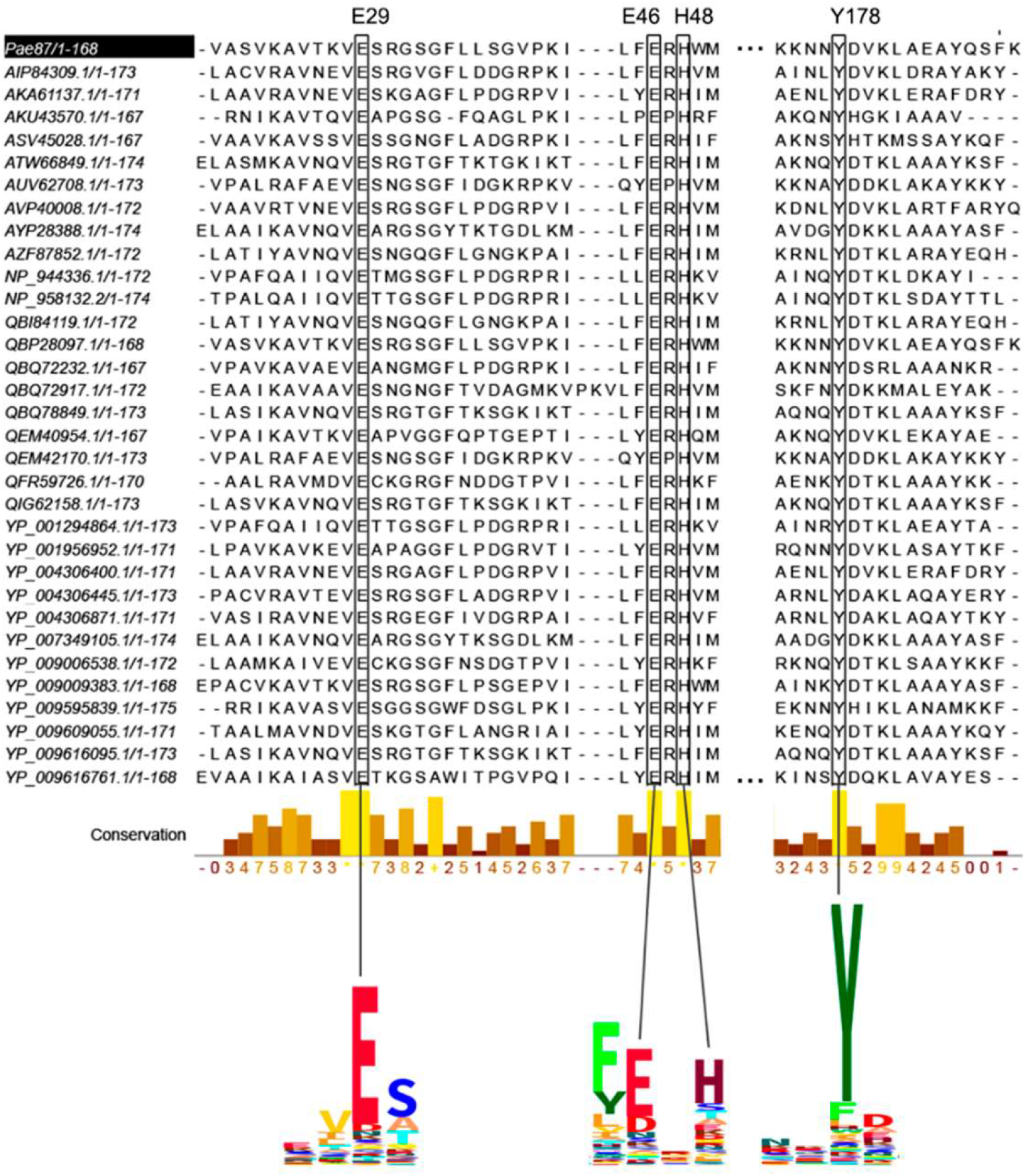
Multiple sequence alignment of *Muramidase* family domains in 𝕊^*MUR*^ showing the conservation of residues facing the catalytic pocket. Selected segments of the PF HMM logo of the family are displayed at the bottom. Relevant positions are connected to their respective columns in the MSA.

No general-base amino acid has been experimentally identified in the *Muramidase* domain family thus far (Maciejewska *et al*., 2017). However, a second Glu residue (Glu46) was found to be perfectly conserved across all of the *Muramidase* examples considered in this work (Fig. 4). This perfect conservation strongly suggests that it plays a relevant role, maybe as the general base, located at a distance of 16 Å from Glu29 (Fig. 2*a*) in a similar fashion as described for the Auto peptidoglycan hydrolase.

Other conserved residues facing the catalytic cleft were His48 and Tyr174, which, therefore, could also play a significant role in the catalytic pocket integrity and function. The aforementioned residues also presented high conservation levels within the PF HMM of the family, judging by the HMM logos (Fig. 4), strengthening their presumptive functional role. In the apo-Pae87 structure, electron density for a CHES buffer molecule was found inside the putative catalytic cleft, in between the putative catalytic glutamates (Fig. 2*cd*). Buffer molecules have also been found in the catalytic cleft of other enzymes, such as the lytic CHAP_K_ domain of the endolysin LysK from *Staphylococcus* bacteriophage K or chitinase C (ChiC) from *Streptomyces griseus* HUT6037 (Sanz-Gaitero *et al*., 2014; Kezuka *et al*., 2006). They may mimic the enzymatic substrate or product, and thus their position reinforces the idea that Glu29 and Glu46 possess catalytic activity in Pae87.

A mutational analysis was conducted to confirm the implication of both Glu29 and Glu46 residues. Pae87 single mutants, E29A and E46A, and a double mutant, E29A/E46A were constructed and their muralytic and bactericidal activities were tested (Fig. 5). Both residues, Glu29 and Glu46, were deemed relevant for the catalytic degradation of PAO1 peptidoglycan since all the mutants displayed a remarkable decrease in their cell wall solubilization ability when compared to wild type Pae87 (Fig. 5*ab*). At the maximum non-saturating concentration (0.1 μM), Pae87 retained about 75% of its maximum activity detected, while the mutants displayed no peptidoglycan solubilization activity. On the other hand, there were no significant differences in the observed bactericidal activity against PAO1 between Pae87 and its mutants or in the fluorescence microscopy images (Fig. 5*cd*).

**Figure 5.**
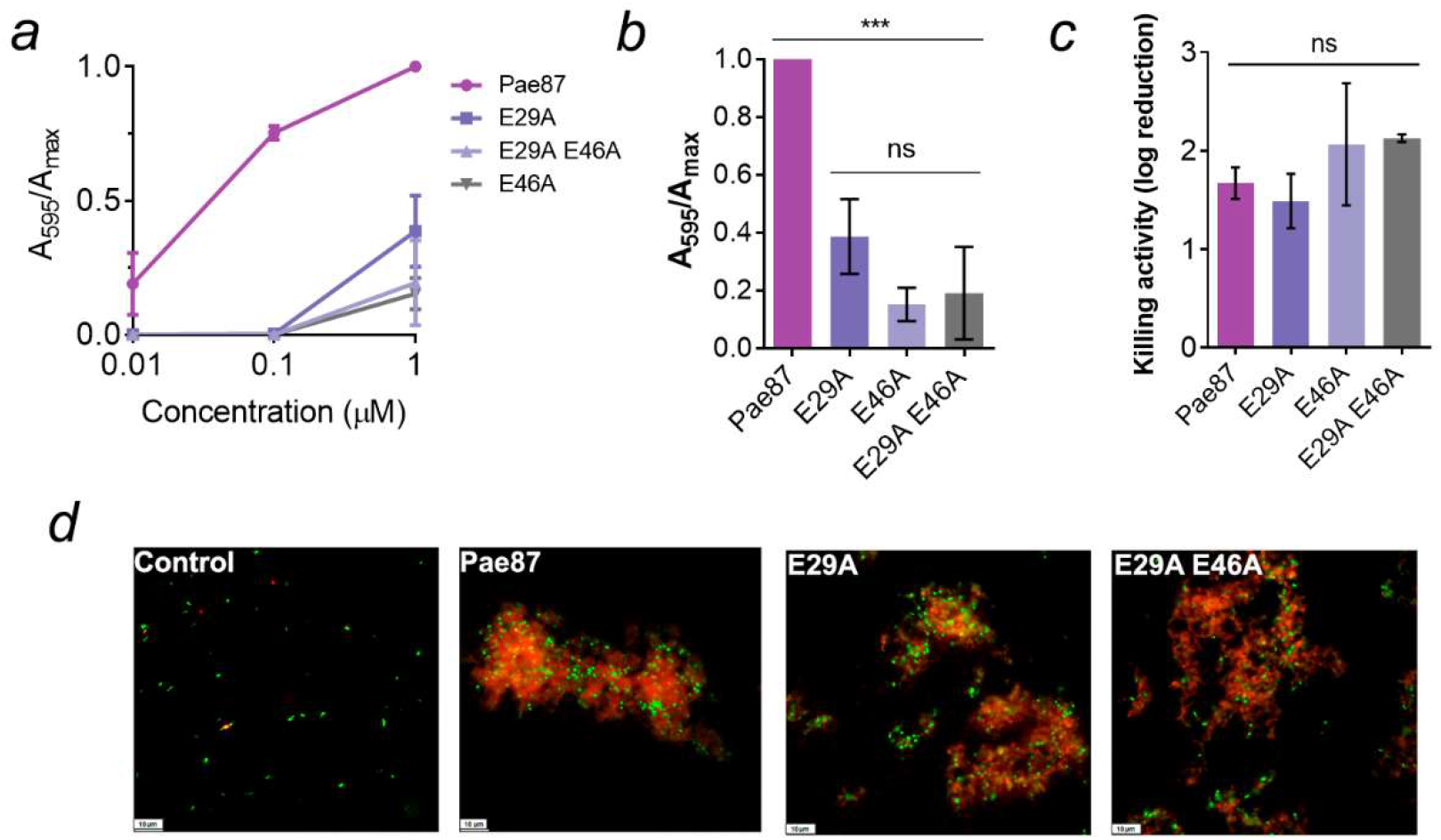
Activities of wild-type Pae87 and of mutants in which conserved catalytic site glutamic acid residues were mutated. (*a*) Muralytic activity of Pae87 and its mutants on RBB-labelled purified PAO1 cell walls at different concentrations. (*b*) Comparison of muralytic activities at the maximum concentration tested (1 μM). (*c*) Bactericidal activity of 10 μM of Pae87 and its mutants against bacterial suspensions of PAO1 (2 h, 37°C). (*d)* Fluorescence microscopy of some of the treated suspensions in (*c*), dyed with SYTO9 and propidium iodide. White bars at the lower-left corner indicate the scale (10 μm). One-way ANOVA was used in (*b*) and (*c*) followed by Tukey post-test to perform an all-against-all multiple comparison (ns = non-significant difference; *** = p ≤ 0.01).

Regarding the catalytic activity of Pae87, the disaccharide found in the soluble product bound into the crystallized protein (NAG-MurNAc, rather than MurNAc-NAG) already points out a lysozyme activity, as described in the literature for this family. The degradation products analysis by RP-HPLC-MS firstly showed that both Pae87 and the positive control —the lysozyme cellosyl (Rau *et al*., 2001)— mobilized an array of soluble compounds, in contrast with the untreated blank (Fig. 6*a*). When comparing the mass spectrometry chromatograms of cellosyl and Pae87, a co-incidental pattern for the main degradation peaks was found (Fig. 6*b*). This observation supports the catalytic nature of Pae87 as a muramidase. Moreover, the CID spectrum for one of the main peaks of Pae87-degraded peptidoglycan (the one eluted at 4.3 min) presents peaks compatible with the loss of a non-reduced NAG (−203.078 mass units) from a NAG-MurNAc-Ala-Glu-mDAP-Ala or a NAG-MurNAc-Ala-Glu-mDAP fragment (Fig. 6*c*). Conversely, there is no evidence coherently compatible with losing a reduced NAG (−223.106).

**Figure 6.**
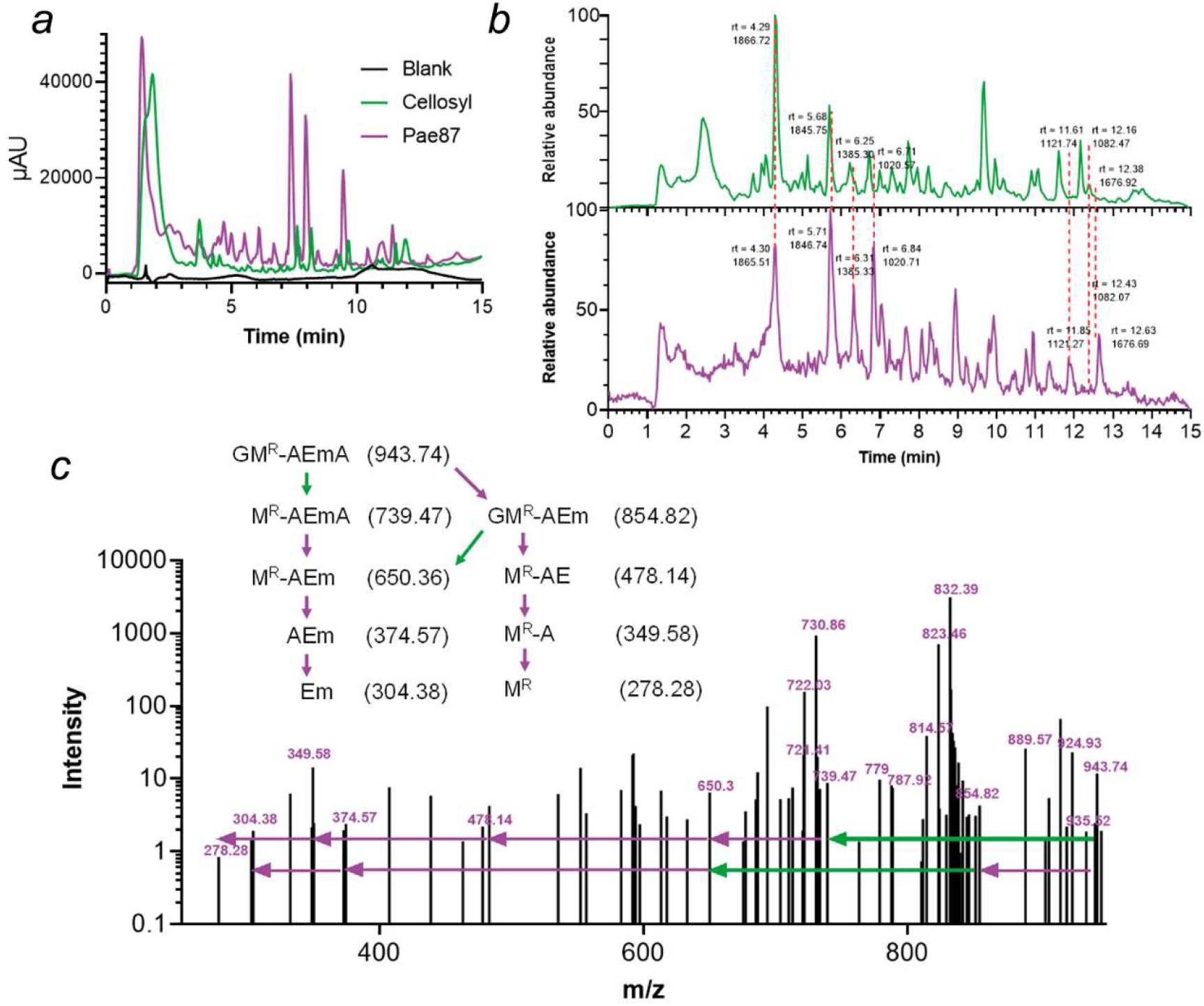
Analysis of the degradation products of Pae87 activity on *P. aeruginosa* PAO1 peptidoglycan. (*a*) UV (204 nm) chromatograms of PAO1 peptidoglycan solubilized with either cellosyl or Pae87 and of the soluble fraction of an untreated sample. (*b*) Liquid chromatography-mass spectrometry chromatograms of the degradation products of cellosyl (green) and Pae87 (purple) activities on PAO1 peptidoglycan. Retention time (rt) and a representative monoisotopic mass value are shown for selected peaks. (*c*) CID spectrum of the rt = 4.30 min peak of the Pae87 peptidoglycan degradation. Selected m/z values are displayed. The dissociation of a GM^R^-AEmA fragment is presented. G = nonreduced NAG; M^R^ = reduced MurNAc; A = alanine; E = glutamic acid; m = meso-diaminopimelic acid.

### 3.4. The noncatalytic activity of Pae87 and P87

The hypothesis that Pae87 displays a noncatalytic bactericidal activity has been previously proposed (Vázquez, Blanco-Gañán, *et al*., 2021), and it is consistent with the results presented in Fig. 5. A closer examination of the physicochemical profile of Pae87 allowed us to postulate a specific 29-amino acids peptide, hereinafter named P87, which would make up a C-terminal AMP-like region (Fig. 7). The definition of P87 was based on the concurrent maximization of net charge and HM that takes place in such coordinates of Pae87. For measuring said properties simultaneously, a score variable was calculated as explained in Materials and methods. The P87 sequence contains seven positively charged Arg or Lys residues (and a negatively charged Glu), interspersed with mostly nonpolar residues. This structure, together with the fact that it forms three alpha-helices within the Pae87 structure (Fig. 7), suggests that it may form amphipathic helices with membrane-interaction potential.

**Figure 7.**
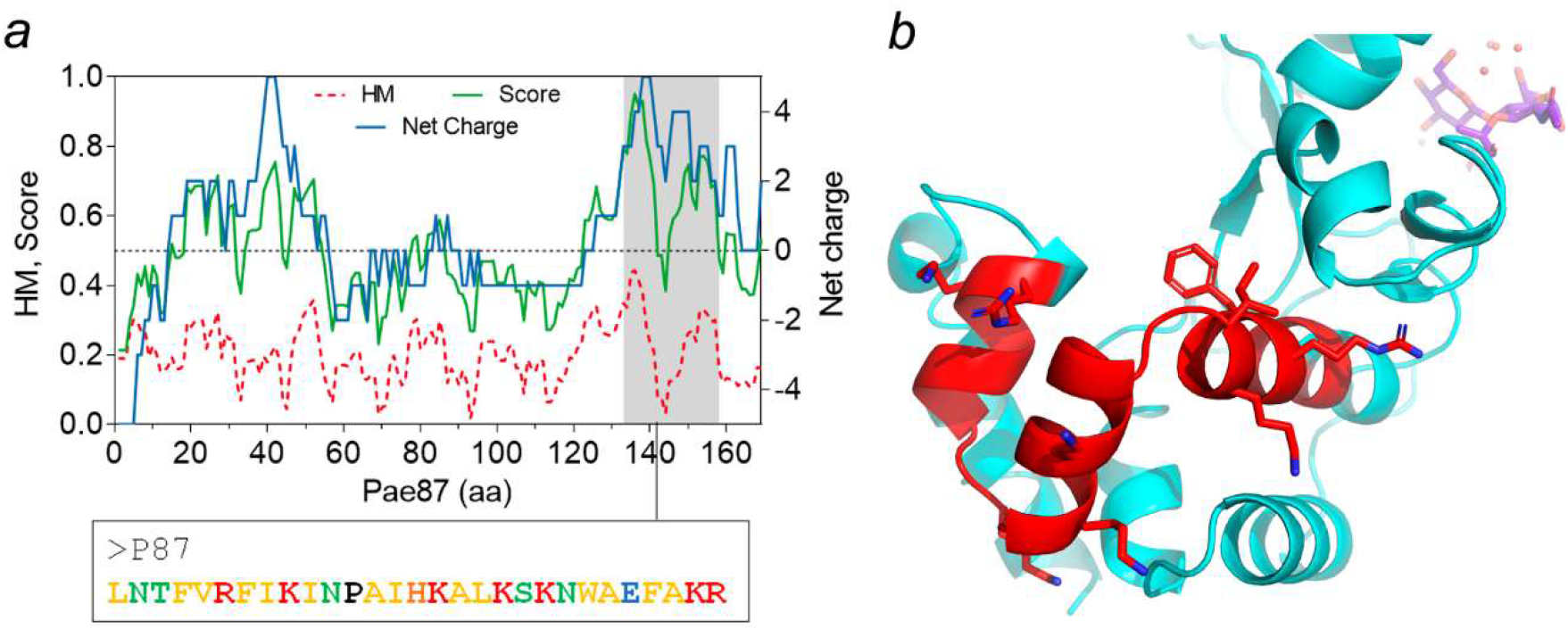
Definition and localization of P87 peptide within Pae87. (*a*) Physicochemical profile of Pae87, depicting HM and net charge in each of the 11-amino acid windows of the protein, as well as the score used to measure both of the properties. P87 sequence is also shown code-coloured regarding the properties of each residue (yellow, nonpolar; green, polar without charge; red, positively charged; black, proline; blue, negatively charged). (*b*) 3D model of Pae87 with peptide P87 highlighted in red. Catalytic Glu residues and peptidoglycan ligand (in blue) are displayed as spatial references.

The synthetic peptide P87 was analyzed for its antimicrobial properties. Circular dichroism spectra in the presence of increasing concentrations of TFE provided evidence on the ability of P87 to form amphipathic helices in the presence of biological membranes (Fig. 8*a*). P87 presented a disordered conformation in an aqueous solution, but, beginning at 20% TFE, it started to shift towards an α-helical conformation, as evidenced by the typical peaks at, roughly, 222 nm and 208 nm. An acute lytic effect of P87 on *P. aeruginosa* PAO1 was observed, in contrast with the lack of generalized lysis when using the full Pae87 protein (Fig. 8*b*). P87 was also able to kill PAO1 and other bacteria in a range and magnitude similar to Pae87 (Fig. 8*c*). The antimicrobial capacity of P87 is thus considered proven, and therefore a role in the Pae87-bacterial surface interaction can be proposed with supporting evidence.

**Figure 8.**
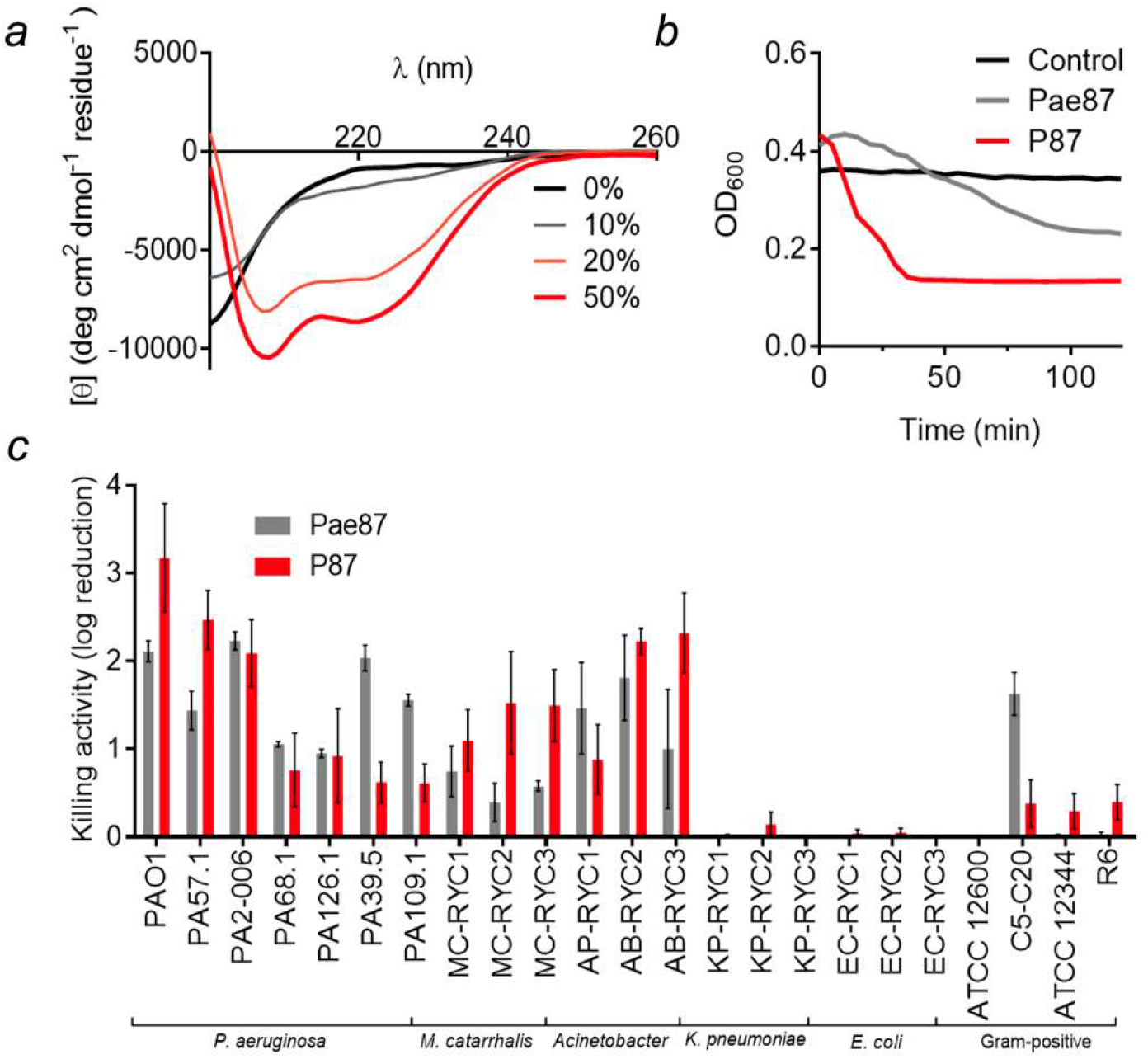
Antimicrobial activity of P87 peptide. (*a*) Far UV circular dichroism spectra of 20 μM P87 in 20 mM NaPiB, pH 6.0, with 100 mM NaCl, and different v/v concentrations of TFE (as indicated in the legend). (*b*) Turbidity decrease assay of a 10 μM P87 or Pae87 treatment to PAO1 cell suspensions. (*c*) Bactericidal activity range of 10 μM P87. Pae87 bactericidal range from (Vázquez, Blanco-Gañán, *et al*., 2021) is also shown for comparison.

### 3.5. Antimicrobial mechanism of Pae87 and P87

Since the OM has traditionally been labeled as the main obstacle when attacking G− bacteria from without, a fluorescent probe was used to detect OM permeabilization induced by Pae87 and its derivatives (*i*.*e*., the noncatalytic mutants and the antimicrobial peptide P87). NPN is a hydrophobic molecule that fluoresces when reaches the phospholipid layer of the inner membrane upon OM permeabilization. Typically, NPN uptake kinetics are recorded with short incubation times (with a maximum of 10 min and a minimum of 3 min) (Loh *et al*., 1984; Helander & Mattila-Sandholm, 2000). However, we extended the incubation period up to 2 h due to the increasing fluorescence values of Pae87 kinetics over such time (Fig. 9*a*). At the average kinetic estimations shown in Fig. 9*a*, two different tendencies can be observed: the ‘canonical’ OM permeabilization peak induced by P87 in the first minutes after peptide addition, and the slow but steady increase in fluorescence for Pae87 and its mutants. For the sake of statistical comparison, the average values of NPN fluorescence reached 5 min after reagent addition were considered (Fig. 9*b*). At such a short period, the NPN fluorescence of Pae87 and its non-catalytic mutants is still low, although, for Pae87, significantly different than the control. The fluorescence induced by P87 was above that obtained for EDTA positive control. Fluorescence microscopy images at the end of incubation visually confirmed the damage to the OM for all the compounds tested (Fig. 9*a*).

**Figure 9.**
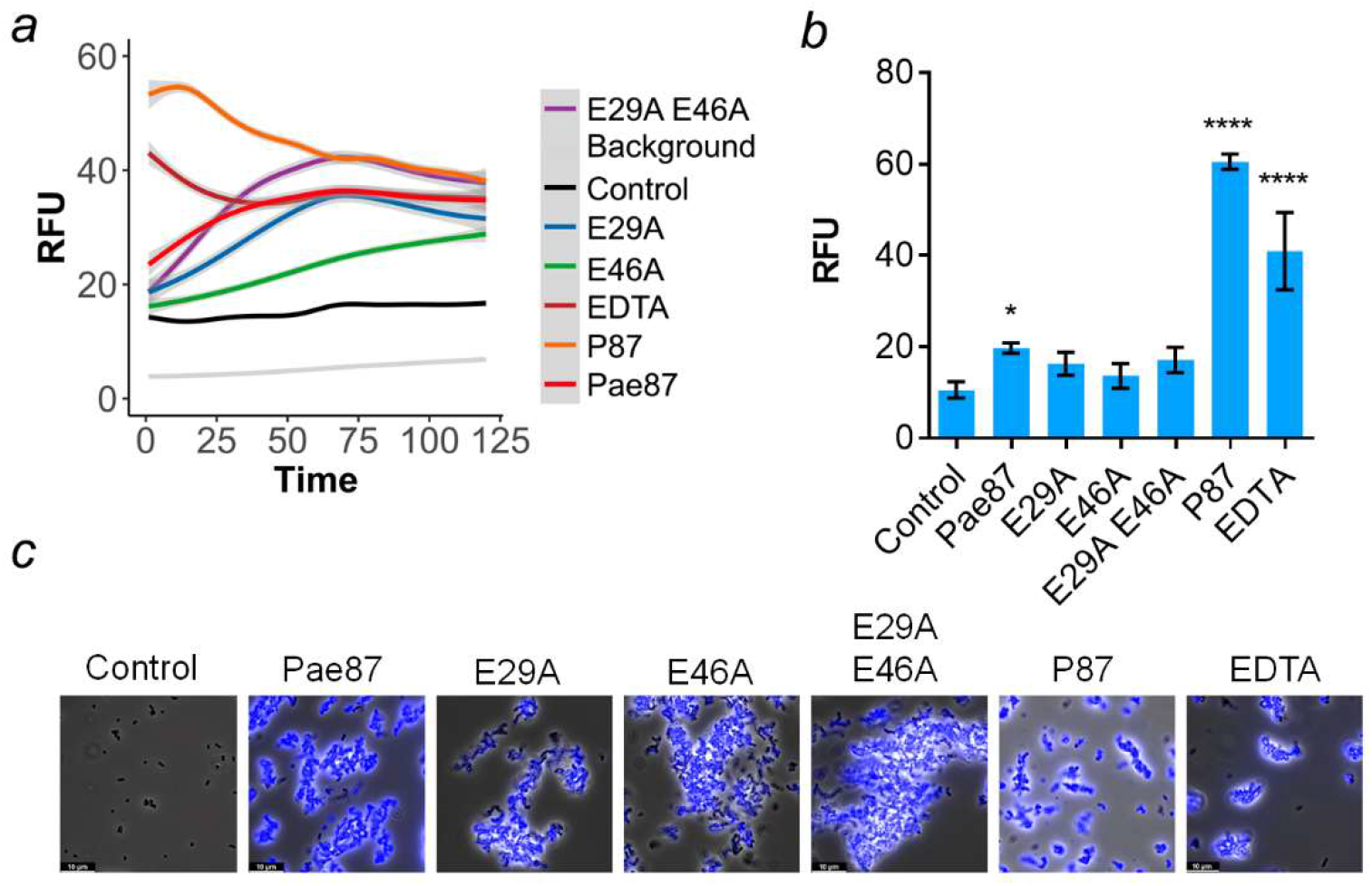
OM permeabilization assays with fluorescent probe NPN. (*a*) GAM estimation of the average tendencies of the NPN fluorescence kinetics (excitation wavelength was 350 nm, emission was recorded at 420 nm). Estimation was based on three independent replicates, mean estimation ± 95% C.I. (grey shade) is shown for each experimental condition. (*b*) Comparison of the average NPN signal (minus background fluorescence) after 5 min incubation. Mean ± sd of three independent replicates is shown; a one-way ANOVA test with Dunnett post-test was applied to statistically compare each condition with the control (untreated cells in the presence of the probe). *, p ≤ 0.05; ****, p ≤ 0.0001. (*c*) Fluorescence microscopy observation of each experimental condition after 2 h incubation. Superimpositions of phase-contrast images with blue fluorescence signal observed with an A filter cube (excitation bandpass 340-380) are shown. RFU = fluorescence units relative to the maximum value achieved during the assay.

A fluorescent version of Pae87 labeled with Alexa488 was used in combination with PI, a DNA-intercalating fluorescent probe. In this way, the temporal localization of Pae87 and its effect on a suspension of PAO1 cells were traced (Fig. 10). At t_0_, Pae87 already began binding to the bacterial surface, as green fluorescence rims were observed around the *P. aeruginosa* cells (Fig. 10*a*). Over time, the Pae87 molecules bound to the cell walls promoted an increasing aggregation among close bacteria. According to the visual estimations of the areas of such aggregates, their maximum size was reached approximately after 30-60 min under the assay conditions (Fig. 10*b*). Also, at t_0_, a discrete intracellular spot of red fluorescence (PI) was observed. Afterwards, PI fluorescence underwent a gradual diffusion concomitant to the aggregation. At 60 min of incubation, red fluorescence appeared as a halo around the aggregates, with a distorted bacterial morphology.

**Figure 10.**
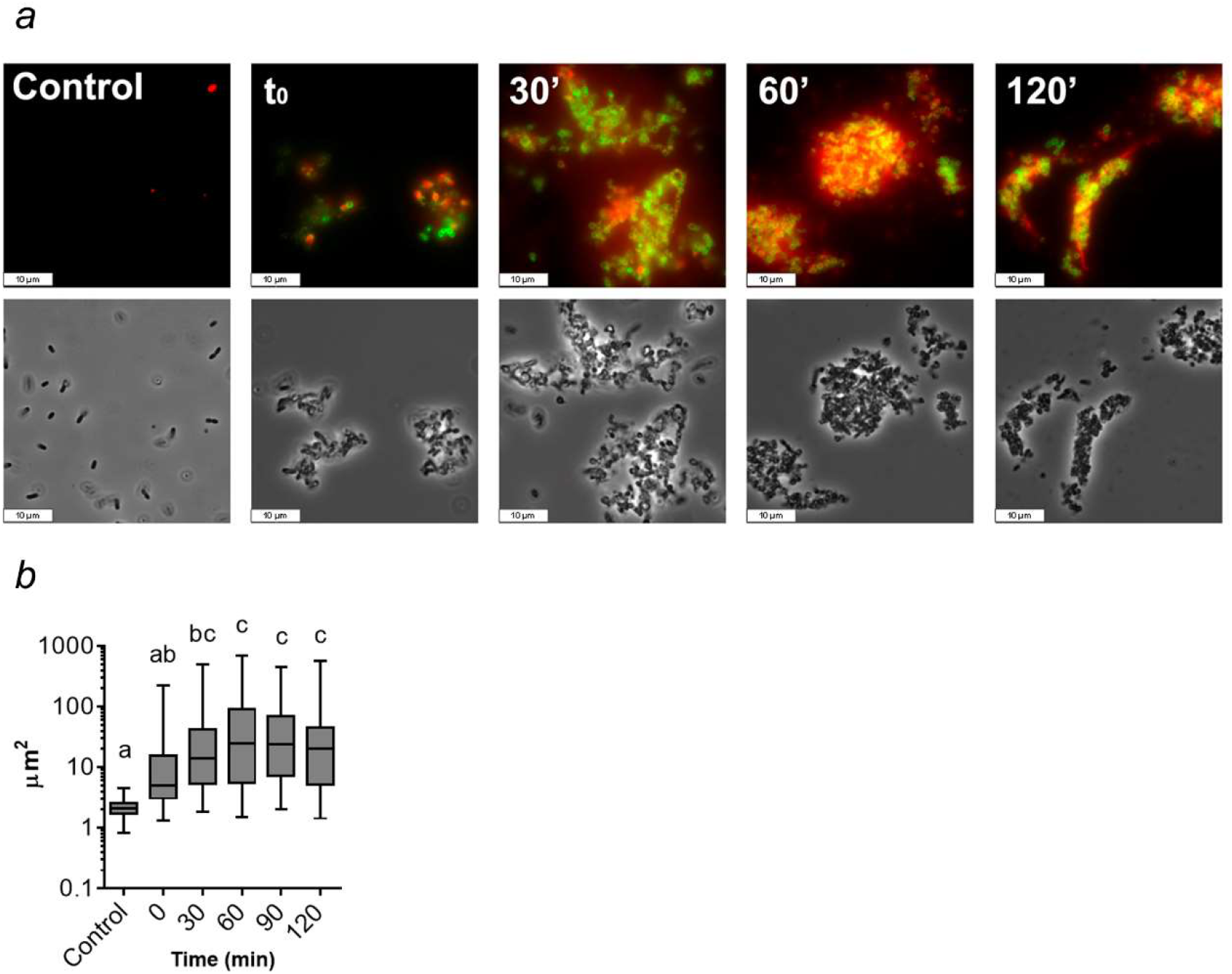
Microscopic observation of a PAO1 suspension treated with Pae87 over time. (*a*) Fluorescence and phase-contrast images representative of the observations made along 2 h of incubation (37°C, 20 mM NaPiB, pH 6.0, 150 mM NaCl) of a PAO1 suspension (≈ 10^8^ CFU ml^-1^) treated with 10 μM of Alexa488-labelled Pae87 and stained with PI. White bars indicate 10 μm. (*b*) Time-wise distributions of cells or aggregates areas estimated using LAS X microscopy image analysis software in, at least, ten frames per time point. A one-way ANOVA followed by Tukey’s post-test was applied for multiple comparisons. Distributions marked with different letters are significantly different from each other, while, between those indicated with the same letter, no significant differences were found.

Several incubation conditions were examined to check their importance in Pae87 and P87 killing activity (Fig. 11). The peptide:bacteria stoichiometry was critical for the killing efficacy of the P87 peptide and, to a lesser extent, for Pae87 (Fig. 11*a*). In both cases, the optimal ratio was achieved at around 10^7^ CFU ml^-1^ for 10 μM Pae87 or P87, which roughly corresponds to some 10^10^ molecules per cell. As for the dose-response curves (Fig. 11*b*), the one of Pae87 quickly saturated, as it was shown before (Vázquez, Blanco-Gañán, *et al*., 2021). This was speculated to be due to the entrapment of Pae87 molecules within the aggregates. P87 presents a more canonical curve, in line with the effect of the peptide:bacteria ratio, with an increasing activity with higher concentrations (up to 5-log kill at the maximum concentration tested).

**Figure 11.**
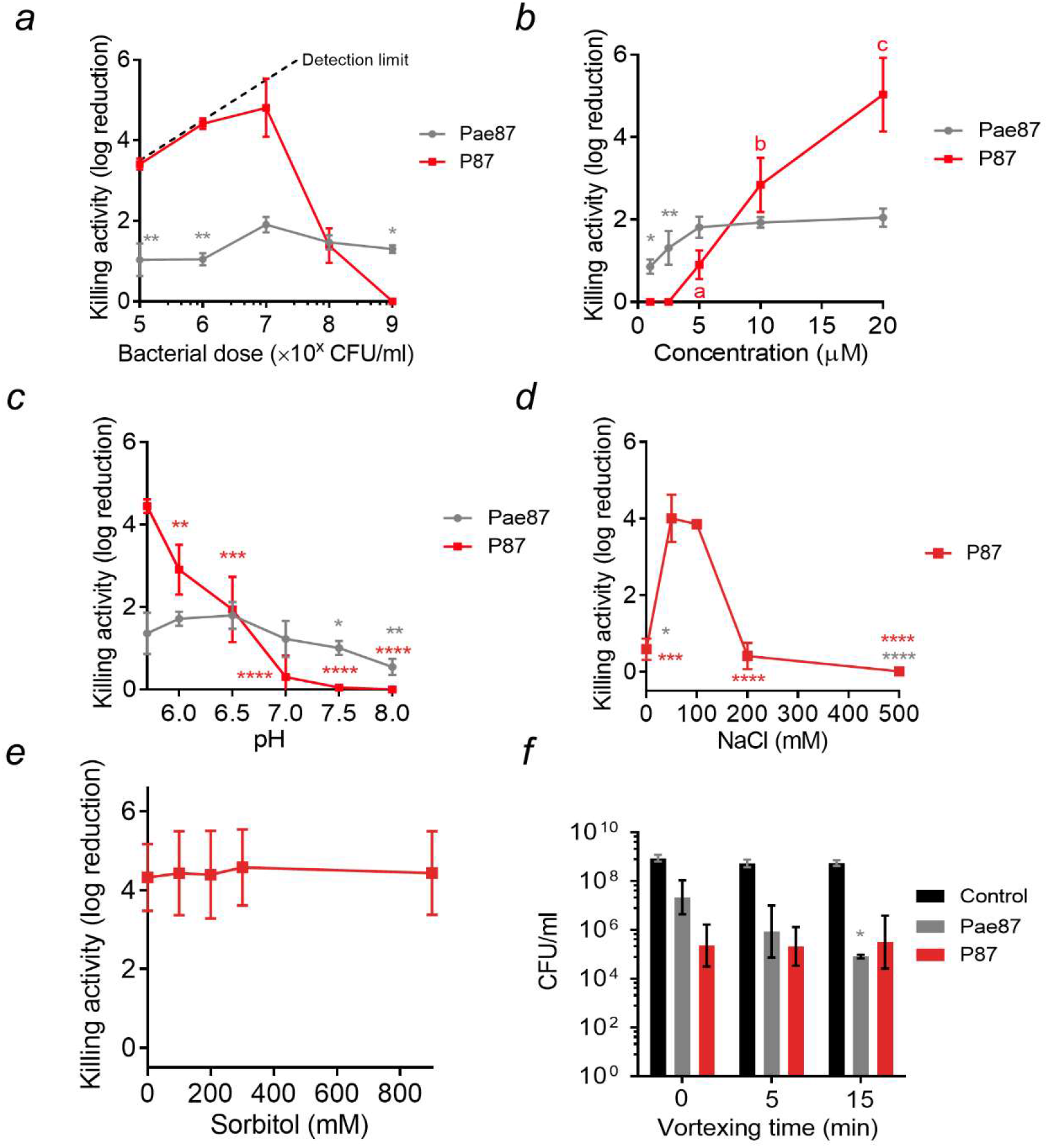
Parameters of Pae87 and P87 bactericidal activity on *P. aeruginosa* PAO1. (*a*) Enzyme:bacteria or peptide:bacteria stoichiometry. (*b*) Dose-response curves. (*c*) Variation of killing activity with pH. (d) Variation of killing activity with ionic strength. (*e*) Variation of killing activity with the concentration of a non-ionic osmolyte (sorbitol), maintaining a fixed concentration of 50 mM NaCl. (*f*) Viable counts after a 0, 5, or 15 min vortexing. One-way ANOVA with Dunnett post-test was applied to compare with the control or the highest value condition (*a*: 10^7^ CFU/ml; *b*: 20 μM; c: 6.5; *d*: 50 or 150 mM; *e*: 0 mM sorbitol; *f*: the corresponding viable count without vortex treatment). Unless otherwise stated, the incubation conditions were 20 mM NaPiB, pH 6.0, 150 mM NaCl, ≈ 10^8^ CFU/ml PAO1, 10 μM of the bactericidal compound, 37°C, 2 h. Asterisks indicate p-values of significant comparisons (* p ≤ 0.05, ** p ≤ 0.01, *** p ≤ 0.001, **** p ≤ 0.0001), non-significant comparisons are not indicated. When marked by letters, an all-against-all multiple comparison was applied with ANOVA plus Tukey post-test. Different letters indicate significantly different results.

pH did not seem a major determinant of Pae87 activity (Fig. 11*c*), although the best killing numbers were obtained at mildly acidic pH, as for P87. Ionic strength did not have a remarkable impact at near-physiological concentrations (50-200 mM NaCl) for Pae87, but P87 only displayed a relevant killing activity between 50 and 150 mM (Fig. 11*d*). It is possible that a slight salt concentration might be necessary for P87 proper solubility (in fact, the Aggrescan server (de Groot *et al*., 2012) detects an N-terminal hotspot for aggregation), but that higher concentrations might shield charged residues. This latter hypothesis is supported by the results in Fig. 11*e*, in which the activity of 10 μM P87 was tested in the presence of different concentrations of a non-ionic osmolyte (sorbitol). In such results, the killing activity of the peptide was not affected by the increasing concentrations of the non-ionic solute. Finally, the 2 h-treated PAO1 suspensions were subjected to different vortexing treatments before plating, to check whether the observed aggregative effect was underestimating the viable cell counts (Fig. 11*f*). The vortex treatment would disintegrate the aggregates and thus release viable cells that, grouped within the aggregates, might have produced a single colony. The results in Fig. 11*f*, however, suggest that this is not the case, *i*.*e*., the observed decrease in viable cell counts cannot be solely attributed to the mere aggregation, and thus a bactericidal mechanism must be occurring. This is because no significant differences were found in the viable counts after vortexing with respect to the non-vortexed samples, except with the samples treated with Pae87 and vortexed for 15 min. In this latter case, the cell count in fact decreased over the control without vortexing.

## 4. Discussion

Knowledge of the structure and function of phage lysins is a crucial step towards a deeper understanding of how lysins work and their application as engineered antimicrobials. Pae87 was previously identified and confirmed as an intrinsically active lysin against a range of Gram-negative pathogens. In this work, such activity has been further investigated in relation to the protein structure to try to unveil the way Pae87 works as an exogenous antimicrobial.

The Pae87 structure shows that it is a one-domain protein with the typical α/β lysozyme fold. In addition, three α-helices are inserted between the β-strands of the β-sheet, forming together the β-lobe. A comparable structure was only found in AP3gp15, a *Burkholderia* phage lysin. A peptidoglycan fragment was bound to this region of the β-lobe in the Pae87-peptidoglycan crystal structure. Interestingly, the residues predicted to be responsible for the peptidoglycan fragment binding were more conserved in representatives from the *Muramidase* family lacking an additional CWBD than in those having it (such as AP3gp15). From these results, the existence of a cell wall binding site within the very catalytic domain can be proposed. In the absence of a CWBD, this region may be used as a binding site by Pae87 and the other related lysins that contain it, perhaps performing a function in the endogenous lytic activity. In fact, most carbohydrate enzymes known to date pose a CWBD that enables them to approach their substrate, since polysaccharidic substrates are insoluble and thus cannot diffuse into the catalytic center (Guillen *et al*., 2010). The observations presented in this work imply that the typically maintained hypothesis of Gram-negative phage lysins lacking or not needing a cell wall binding function (Ghose & Euler, 2020; Vázquez, García, *et al*., 2021) should be perhaps revised once sufficient evidence is gathered on i) whether these internal regions within the catalytic domain function as true CWBDs and ii) how widespread this trait is. To our knowledge, there are few examples in literature of cell wall binding sites contained within the catalytic domain. For example, *Bacillus* lysin PlyG contains a region located within the NAM-amidase catalytic domain that specifically recognizes *Bacillus anthracis* spores (Yang *et al*., 2012). From a functional point of view, this is hardly a comparable case, since the secondary substrate binding site of PlyG seems to have evolved to recognize a chemically distinct form of the target bacteria that cannot be recognized by the canonical CWBD. A binding region far from the active site has been also proposed at the C-terminal lobe of T4 lysozyme (Grutter & Matthews, 1982; Kuroki *et al*., 1993), in this case proposing that it binds the peptide moiety of the peptidoglycan thus providing some specificity to the lytic activity. In contrast, for Pae87 and its close relatives, it could be argued that a region with affinity to peptidoglycan has arisen to take over the function of a lacking CWBD, given the differential conservation of the predicted key residues for binding.

Unlike the binding motif, the Pae87 residues of the catalytic cleft are conserved in AP3gp15 (Fig. 2*b*). Here, the most important residues are Glu29 and Glu46, responsible for the catalytic activity. Glu29 may correspond to the general acid of other enzymes of the lysozyme family (Wohlkonig *et al*., 2010). An equivalent to Glu46, the presumed general base, is only found in some lysozymes, such as the peptidoglycan hydrolase Auto (Bublitz *et al*., 2009) and AP3gp15, although the function of the Glu46 equivalent in the highly homologous AP3gp15 was not studied and, in fact, it was not pointed out as a possible catalytic residue (Maciejewska *et al*., 2017). The conserved residues at the MSAs presented in this work have allowed the identification of said glutamic acid and we also showed that the alanine mutant of Glu46 abolishes lysozyme activity in Pae87, therefore confirming the catalytic function of both Glu29 and Glu46. However, the distance between the glutamates is 16 Å, longer than the typical distance of inverting glycoside hydrolases (Davies & Henrissat, 1995). For context, the typical O…O inter-carboxylic distance has been recorded to be, in average, 8.5 ± 2.0 Å in inverting β-glycosidases, while it is shorter (4.8 ± 0.3 Å or 6.4 ± 0.6 Å) in β-glycosidases that use the retaining mechanism (Mhlongo *et al*., 2014). A putative conclusion from these observations is that the hydrolysis mechanism of Pae87 may operate with net inversion of the anomeric configuration. In fact, Bublitz and collaborators proposed Auto could move to a closed conformation when it binds to its substrate and follow the inverting mechanism of glycoside hydrolysis. The same conformational readjustment and mechanism could occur in Pae87. While mutating Glu29 and Glu46 dramatically diminished Pae87 muralytic efficacy, the three mutants (including E29/E46 double mutant) also retained some residual activity at the maximum concentration tested (less than 50% than the wild type enzyme), suggesting that there may be other residues involved in catalysis (perhaps such conserved residues as His48 and/or Tyr174), or rather that some residues in the catalytic cleft may take over the function of the mutated amino acid to some degree. The previous analyses on the precise catalytic activity of lysins from the *Muramidase* family have all concluded that it actually poses a muramidase or lysozyme activity that breaks the glycan chain of the peptidoglycan on the reducing side of MurNAc (Rodriguez-Rubio *et al*., 2016; Maciejewska *et al*., 2017). Our results also fit with this proposal. To begin with, correspondence was found between the main degradation peaks detected by RP-HPLC-MS when treating *P. aeruginosa* peptidoglycan with cellosyl, a known lysozyme, and Pae87. Some of the coincidental main peaks can be easily assigned to peptidoglycan degradation products. For example, the main peaks with monoisotopic m/z ≈ 1865 or 1845 are consistent with fragments containing two disaccharides (NAG-[reduced MurNAc]) crosslinked by two tetrapeptides (Ala-Glu-mDAP-Ala). The difference in mass (1865 vs 1845) corresponds to losing a water molecule, probably in one of the reduced MurNAc residues. The CID results are definitive proof for a muramidase activity, which leaves a MurNAc end susceptible to be reduced (conversely, NAG is susceptible to reduction when the glycan strand is degraded by a glucosaminidase activity) (Eckert *et al*., 2006; Rodriguez-Rubio *et al*., 2016).

On another hand, the lack of a difference in bactericidal activity between Pae87 and its mutants suggests that the membrane activity is the major determinant for Pae87 antimicrobial potential, rather than the catalytic activity, as pointed out for other intrinsically active lysins (Ibrahim *et al*., 1996). In this work, a specific AMP-like C-terminal region (P87) with intrinsic membrane-permeabilizing and bactericidal activity has been identified as the most probable part of the enzyme responsible for the aforementioned effect. The crystal structure confirmed that P87 was located at the surface of the protein, supporting the hypothesis that it would be able to directly interact with membranes. Moreover, Pae87 had an OM permeabilizing activity by itself as demonstrated by the NPN uptake assay. Nonpolar residues of P87 are mostly buried within Pae87, but not in all cases. For example, Ile164 or even Phe161 are especially exposed (Fig. 7*b*). Lys and Arg residues of P87, on the other hand, are located at the outer surface of the protein, therefore available for electrostatic interaction with negatively charged elements of the bacterial surface (namely, the phosphate groups of the lipopolysaccharide). Based on the results in this work, including the micrographs presented in Fig. 10, a mechanism for Pae87 activity from without is proposed: i) the P87 region of the enzyme would bind to the OM, coating the bacterial surface and then causing cause the aggregation of adjacent cells; ii) then, the membrane-permeabilizing action would act, perhaps together with the peptidoglycan hydrolysis activity, to disrupt the cell wall; iii) the leakage of intracellular components and cell death takes place without provoking a full disintegration of the bacteria (‘lysis’), but rather keeping the cell debris tightly bound in compact aggregates. The viability decreases results after vortexing presented in Fig. 11 are in agreement with the proposal that aggregates comprise damaged cells (not relatively intact ones) and, thus, the mechanical shaking increased the apparent killing by definitely harming these already damaged bacteria. This ‘death without lysis’ could be beneficial from the point of view of *in vivo* therapy, since it would prevent the dissemination of pro-inflammatory factors. Also, bacterial aggregates have been previously shown to be better cleared by the immune system (Ribes *et al*., 2013; Roig-Molina *et al*., 2020). However, the true potential of this kind of antimicrobial agent should be tested *in vivo* to clarify its possible benefits.

Regarding the antimicrobial activity of peptide P87 itself, it was observed to cause an acute lytic effect on *P. aeruginosa* cells. The ability to form amphipathic helices in the presence of TFE, proved by the circular dichroism spectra in Fig. 8, would point to the insertion of such helices into the biological membranes and subsequent leakage of the cell contents as the mechanism for P87-mediated killing. The peptide differs, in this regard, with the poor lytic outcome of Pae87 treatment. This difference may be due to the much smaller size of P87, which could enable it to properly insert into the membranes rather than interact superficially, as it is assumed that Pae87 does.

The maximum detectable killing was observed for 10 μM P87 at a cell concentration of 10^7^ CFU ml^-1^ and below, while higher bacterial doses reduced the bactericidal effect, surely due to a sub-optimal number of antimicrobial molecules per cell. This is in agreement with a cooperative mechanism of action where a threshold number of bound peptides is required for the bactericidal activity. In addition, the higher activity observed at more acidic pHs can be explained due to the higher positive charge that both protein and peptide may have at acidic pHs, improving interaction with the negatively charged bacterial surface. Given that the activity increase was observed at pH 6.0 and below, the protonation of histidine residues (pK_a_ ≈ 6.0) is the most plausible explanation. Although P87 was almost inactive at near-physiological pH (≈ 7.5), the fact that it was highly effective at acidic pH is relevant for infection treatment, since it has been many times suggested that the pH at the infection site is acidified by a combination of bacterial metabolic activity and immune system responses (Radovic-Moreno *et al*., 2012; Simmen & Blaser, 1993). This acidification is especially relevant in certain conditions, such as cystic fibrosis, whose patients are already infection-prone at the respiratory tract, with *P. aeruginosa* being one of the main causative agents of cystic fibrosis exacerbation (Poschet *et al*., 2002). The importance of charge is also manifested by the ionic strength experiments: P87 only displayed a relevant killing activity between 50 and 150 mM NaCl. It is possible that a slight salt concentration might be necessary for P87 proper solubility. Higher concentrations might, however, shield charged residues. This latter hypothesis is supported by the results in the presence of different concentrations of a non-ionic osmolyte (sorbitol), in which the killing activity of the peptide was not affected by the increasing concentrations of the solute.

## Conclusions

The three-dimensional structure of Pae87 has been elucidated by X-ray crystallography. This structure provided the basis to propose the presence of a substrate-binding subdomain within the catalytic domain of Pae87. This substrate-binding site is apparently conserved among other enzymes from the same family that lack an independent CWBD and thus may fulfil a compensatory evolutionary function. It was determined that Pae87 is a muramidase, and two acidic residues have been pointed out as involved in such catalytic activity. However, the antimicrobial activity of Pae87, when exogenously added, was not associated with the catalytic activity, but rather to a nonenzymatic activity on the membranes that most probably resides on a cationic, amphiphilic C-terminal peptide named P87. Such a peptide was proven to be an AMP on its own, active against a range of bacteria coincidental to those susceptible to Pae87 surface activity. The activity of P87 was highly dependent on its intrinsic charge, and on the peptide:bacteria stoichiometry. Altogether, these results provide further clarity on the intra-structure of a family of Gram-negative-active lysins, revealing evolutionary features with a close relationship to architectural traits of the bacterial hosts. On the other hand, some insights have also been provided on the intrinsic antibacterial effect of Pae87 and a novel AMP, P87, has been discovered.

## Acknowledgements

The ALBA-CELLS beamline BM13 (XALOC) and Fernando Gil and Xavier Carpena are thanked for providing excellent crystallographic data collection facilities.

## Funding information

The authors acknowledge the Spanish Ministry of Science and Innovation for grant numbers SAF2017-88664-R (to PG) and BFU2017-87022-P (to MJvR) funded by MCIN/AEI/10.13039/501100011033 and by ERDF A way of making Europe, and for the Severo Ochoa program to the CNB-CSIC (SEV 2017-0712). Additional funding was provided by the Centro de Investigación Biomédica en Red de Enfermedades Respiratorias (CIBERES) to PG, an initiative of the Instituto de Salud Carlos III. RV was the recipient of a predoctoral fellowship from CIBERES.

## Supporting information

**Table S1**. Plasmids and oligonucleotides used throughout this work.

**Table S2**. Protein parameters as predicted by ProtParam

## Notes

### Competing Interest Statement

The authors have declared no competing interest.

