## Supporting Information for "Monomodular *Pseudomonas aeruginosa* phage JG004 lysozyme (Pae87) contains a bacterial surface-active antimicrobial peptide-like region and a possible substrate-binding subdomain"

^d^Interdisciplinary Platform for Sustainable Plastics towards a Circular Economy (SusPlast-CSIC), Madrid (Spain).

^1^These authors contributed equally

**Table S1**. Plasmids and oligonucleotides used throughout this work.

| **Name** | **Description** | **Reference or source** |
| --- | --- | --- |
| Plasmids | | |
| pET-PA87 | Derived from pET-28a(+) and pUCPA87, overexpresses gene *pae87* for production of protein Pae87, fused to a 6×His tag at N-terminal end. KAN^R^ | (Vázquez *et al.*, 2021) |
| pET-PA87-E29 | Derived from pET-PA87, overexpresses gene *pae87-e29* which encodes protein Pae87 with mutation E29A. KAN^R^ | This work |
| pET-PA87-E46 | Derived from pET-PA87, overexpresses gene *pae87-e46* which encodes protein Pae87 with mutation E46A. KAN^R^ | This work |
| pET-PA87-E2946 | Derived from pET-PA87-E29, overexpresses gene *pae87-e2946* which encodes protein Pae87 with mutations E29A and E46A. KAN^R^ | This work |
| Oligonucleotides (5’ 🡪 3’) | | |
| pae87_f | CTAAGGTACCCATATGGCTCTGACCGAGCAAGACTTCC | 5' of *pae87* (forward) |
| pae87_3' | TACAAAGCTTATTTGAAGGATTGATAGGCTTCTGCCAG | 3' of *pae87* (reverse) |
| e29a_f | CGTCACCAAAGTAGCGAGTCGTGGG | Triplet coding for E29 of *pae87*, for mutation (forward) |
| e29a_r | CCCACGACTCGCTACTTTGGTGACG | Triplet coding for E29 of *pae87*, for mutation (reverse) |
| e46a_f | TTCTGTTCGCACGCCACTGG | Triplet coding for E46 of *pae87*, for mutation (forward) |
| e46a_r | TGGCGTGCGAACAGAATTTTCGG | Triplet coding for E46 of *pae87*, for mutation (reverse) |

Restriction enzyme recognition sites are underlined and mutated bases with respect to the wild-type sequence are highlighted in grey.

**Table S2**. Protein parameters as predicted by ProtParam (Artimo *et al.*, 2012).

| **Protein** | **Molecular mass (kDa)** | **Number of aa** | **p*I*** | **Molar extinction coefficient (M^-1^ cm^-1^)** |
| --- | --- | --- | --- | --- |
| Pae87 | 23.05 | 206 | 9.11 | 32555 |
| E29A | 22.99 | 206 | 9.24 | 32555 |
| E46A | 22.99 | 206 | 9.24 | 32555 |
| E29A E46A | 22.93 | 206 | 9.35 | 32555 |

Artimo, P., Jonnalagedda, M., Arnold, K., Baratin, D., Csardi, G., de Castro, E., Duvaud, S., Flegel, V., Fortier, A., Gasteiger, E., Grosdidier, A., Hernandez, C., Ioannidis, V., Kuznetsov, D., Liechti, R., Moretti, S., Mostaguir, K., Redaschi, N., Rossier, G., Xenarios, I. & Stockinger, H. (2012). *Nucleic Acids Res* **40**, W597-603.

Vázquez, R., Blanco-Gañán, S., Ruiz, S. & García, P. (2021). *Front Microbiol* **12**, 660403.
